# Caspase-2 kills cells with extra centrosomes

**DOI:** 10.1101/2024.02.13.580097

**Authors:** Dario Rizzotto, Vincenza Vigorito, Patricia Rieder, Filip Gallob, Gian Mario Moretta, Claudia Soratroi, Joel S. Riley, Florian Bellutti, Stefano Li Veli, Alessia Mattivi, Michael Lohmüller, Sebastian Herzog, Beat C. Bornhauser, Etienne D. Jacotot, Andreas Villunger, Luca L. Fava

## Abstract

Centrosomes are membrane-less organelles that orchestrate a wide array of biological functions by acting as microtubule organizing centers. Here, we report that caspase-2-driven apoptosis is elicited in blood cells failing cytokinesis and that extra centrosomes are necessary to trigger this cell death. Activation of caspase-2 depends on the PIDDosome multi-protein complex and priming of PIDD1 at extra centrosomes is necessary for this pathway. Accordingly, loss of its centrosomal adapter, ANKRD26, allows for cell survival and unrestricted polyploidization in response to cytokinesis failure. Mechanistically, cell death is initiated upstream of mitochondria via caspase-2-mediated processing of the BCL2 family protein BID, driving BAX/BAK-dependent mitochondrial outer membrane permeabilization (MOMP). Remarkably, BID-deficient cells enforce apoptosis by engaging p53-dependent pro-apoptotic transcriptional responses initiated by caspase-2. Consistently, BID and MDM2 act as shared caspase-2 substrates, with BID being kinetically favored. Our findings document that the centrosome limits its own unscheduled duplication by the induction of PIDDosome-driven mitochondrial apoptosis to avoid potentially pathogenic polyploidization events.

## Introduction

Centrosomes are membrane-less organelles involved in multiple biological processes, including mitotic spindle pole organization, cell migration, immune-synapse formation, as well as ciliogenesis (*1*, *2*). Lack of centrosomes has been causally linked to defective neurodevelopment and neurological disorders due to loss of neuronal progenitor cells (*3*), a condition recapitulating mutation of structural components of the centrosome found in human microcephaly (*4*). Excess centrosomes instead can impair faithful microtubule-kinetochore attachments in pseudo-bipolar mitotic spindles or promote multi-polar mitoses, thereby increasing the risk of cells acquiring an aneuploid karyotype (*5*, *6*). Unsurprisingly, centrosome abnormalities are frequently found in cancer and forced centrosome overduplication by PLK4 overexpression, a key regulator of their biogenesis, has been reported to suffice to promote cancer formation in animal models (*7*). Whether extra centrosomes are cause or consequence of malignant transformation is still debated, as healthy cells respond with p53-induced cell cycle arrest to their accumulation (*8*). Consistently, loss of p53 facilitates proliferation of immortalized cells overexpressing regulators of centriole duplication (*9*). These observations firmly establish p53 as a barrier against aneuploidy and transformation resulting from extra centrosomes. Whether this barrier is imposed solely by p53-dependent cell cycle arrest or whether it involves additional effector mechanisms remains uncertain.

Caspases are a class of endopeptidases that are recognized arbiters of cell death and inflammation. Caspase-2 is the most conserved member of the caspase-family and is activated in the PIDDosome multi-protein complex (*10*, *11*). Its overall role in apoptosis, however, has remained a matter of intense debate. Triggers reported to promote caspase-2 activation include protein aggregation upon heat shock, ER-stress, chronic spindle assembly checkpoint (SAC) activation, DNA damage, as well as G2/M checkpoint override (*12–15*). However, studies employing multiple cell types from animals lacking individual PIDDosome components, including PIDD1 and RAIDD, failed to provide support for a rate-limiting role of caspase-2 to cell death initiation in response to this broad array of triggers (*16–18*). Regardless of these discrepancies, caspase-2 has been assigned tumor suppressor potential in animal models of *MMTV/c-neu*-driven breast cancer, ATM-loss and MYC-driven lymphomagenesis (*19–21*). Of note, in the latter, it was suggested to act by deleting aneuploid cells, as mouse embryonic fibroblasts and MYC-driven B cell lymphomas lacking caspase-2 showed increased frequencies of abnormal karyotypes (*22*, *23*). Moreover, a large-scale clinical study documented reduced caspase-2 expression levels to correlate with increased aneuploidy tolerance in colorectal cancer patients lacking the WNT pathway modulator, BCL2L9 (*24*). Consistent with the idea that caspase-2 can kill aneuploid cells, loss of caspase-2 in HCT116 colon cancer cells provided partial protection from cell death induced by the aneuploidy-inducing reagent Reversine, an inhibitor of MPS1 kinase crucial for spindle assembly checkpoint activity (*24*). However, Reversine also induces cytokinesis failure in a significant fraction of cells traversing mitosis in the absence of a functional SAC (*8*). Taken together, these observations suggest that caspase-2 has a role in limiting the survival or the proliferative potential of aneuploid cells. Yet, it still remains unclear whether the activating cue for caspase-2 resides in the aneuploidy itself or rather a feature arising concomitantly to aneuploidy, such as extra centrosomes. Moreover, caspase-2 effectors promoting cell death are poorly defined. The pro-apoptotic BCL2 family protein BID, a member of the BH3-only subgroup, has been reported as a caspase-2 substrate and implicated in subsequent cell death initiation. Yet, most triggers reported to engage caspase-2 for apoptosis induction, such as DNA damage or ER stress do not seem to universally rely on BID being present (*25*, *26*). BID plays a well-established role in connecting the intrinsic mitochondrial and extrinsic apoptosis pathways in response to death receptor activation. For this function it needs to be processed by caspase-8 into its active truncated form, tBID (*27*). However, a physiological trigger driving processing of BID in a caspase-2-dependent manner remains to be defined. P53 activation in response to centrosome accumulation requires assembly of the PIDDosome, a multi-protein complex able to activate pro-caspase-2. While early studies placed the PIDDosome downstream of p53 in the context of DNA damage-induced apoptosis, with PIDD1 being a transcriptional p53 target (*11*, *28*), our recent findings document that it can also act upstream by promoting caspase-2-dependent proteolysis of MDM2, leading to p53 accumulation in cells that accumulate extra centrosomes (*8*). Of note, the ability of the PIDDosome to respond to extra centrosomes is tightly linked to the presence of extra mature parent centrioles. Mechanistically, physical association of PIDD1 with mature parent centrioles requires centriolar distal appendages and ANKRD26 as connecting element (*29*, *30*). Similar to the loss of one of the three PIDDosome components, PIDD1, RAIDD or caspase-2, ANKRD26 deficiency abrogates MDM2 cleavage and p53 pathway activation. This response is engaged in cancerous and immortalized epithelial cell lines that accumulate centrosomes, as well as in primary hepatocytes that undergo scheduled cytokinesis failure during development or regeneration (*8*, *31*). Curiously, in tissue culture, cells carrying extra centrosomes lose them over time due to strong counterselection (*6*, *32*). Whether this phenomenon is solely due to impaired proliferation of cells that fail to normalize their centrosome number or active promotion of cell death downstream of centrosome accumulation remains uncertain.

Considering the purported role of caspase-2 in promoting apoptosis of aneuploid cells as opposed to promoting cell cycle arrest in cells accumulating centrosomes, it is presently unclear whether different cues lead to different outcomes of caspase-2 activation, or rather cell type and context drive fate switch once caspase-2 becomes activated by a common trigger. Here, we provide evidence that caspase-2 promotes apoptosis downstream of extra centrosomes by targeted proteolysis of its shared substrates, BID and MDM2. Our findings reconcile a vast set of seemingly contradictory reports in the field on the apoptotic contribution of caspase-2 and provide a rationale for the activity of a new class of anti-mitotic agents targeting the SAC and mitotic progression, such as Aurora kinase, MPS1 kinase or CENP-E inhibitors.

## Results

### Mitotic perturbation leads to variable outcomes depending on the cell type

Initially, we analyzed the response of different cell types with functional p53 status to different inhibitors of mitotic progression or cytokinesis, reported to promote p53 activation. These included the spindle poisons Nocodazole (Noc) or Taxol (Tax), which activate the spindle assembly checkpoint. We also used a range of inhibitors which promote cytokinesis failure, including dihydrochalasin B (DHCB), the Aurora kinase inhibitor, ZM447439 (ZM), or a combination treatment of Taxol + Reversine (Tax+Rev), an MPS1 kinase inhibitor. Reversine, overriding the SAC, forces cells to rapidly progress through mitosis irrespectively of the presence of unattached kinetochores. We exposed human A549 lung cancer cells, the immortalized hTERT-RPE1 retinal pigmented epithelial (RPE1) cells and Nalm6 pre–B ALL, as well as murine BaF3 pro-B cells to the aforementioned treatments (Fig. S1A). While Nocodazole did not induce significant death in epithelial cells, Taxol did so quite potently in all the cell lines tested, epithelial and hematopoietic alike. However, we also noted that epithelial cell lines consistently entered cell cycle arrest after treatment with cytokinesis failure-inducing drugs, while hematopoietic cells readily undergo apoptosis (Fig. S1B).

### An unbiased genetic screen identifies the PIDDosome as an activator of mitochondrial apoptosis in cells that fail cytokinesis

To identify regulators and effectors of cell death induced by SAC activation or cytokinesis failure, we employed an unbiased forward genetic screen using a genome-wide sgRNA CRISPR lentiviral library. Taking advantage of the sensitivity of BaF3 cells to cytokinesis failure-induced cell death, we transduced a Cas9-expressing derivative clone with a sgRNA library and exposed them to Taxol or Taxol in combination with Reversine for 48 hours (Fig. 1,A and B). Surviving cells were isolated by Ficoll-gradient purification and the enriched sgRNAs were profiled by next generation sequencing (NGS). Using a cut-off of at least 3 out of 6 sgRNA per gene with a p-value < 0.05, we identified a series of established genes known to promote mitochondrial apoptosis in cells after Taxol treatment. In addition to p53, sgRNAs targeting *Bim/Bcl2l11, Bbc3/Puma, Dffb/Cad, Itpr2, Apaf-1 and Casp9* were enriched. Interestingly, in the Taxol + Reversine condition, we similarly found apoptotic regulators (*Trp53*, *Bbc3*, *Bax*, *Bim*, *Apaf-1*, *Casp-9*), as well as key regulators of the centrosome-PIDDosome signaling axis, notably *Cep83/Ccdc41, Ankrd26,* in addition to the PIDDosome components *Pidd1/Lrdd, Raidd/Cradd* and *Casp2* (Fig. 1,C and D and Fig. S2, A and B).

**Fig. 1:**
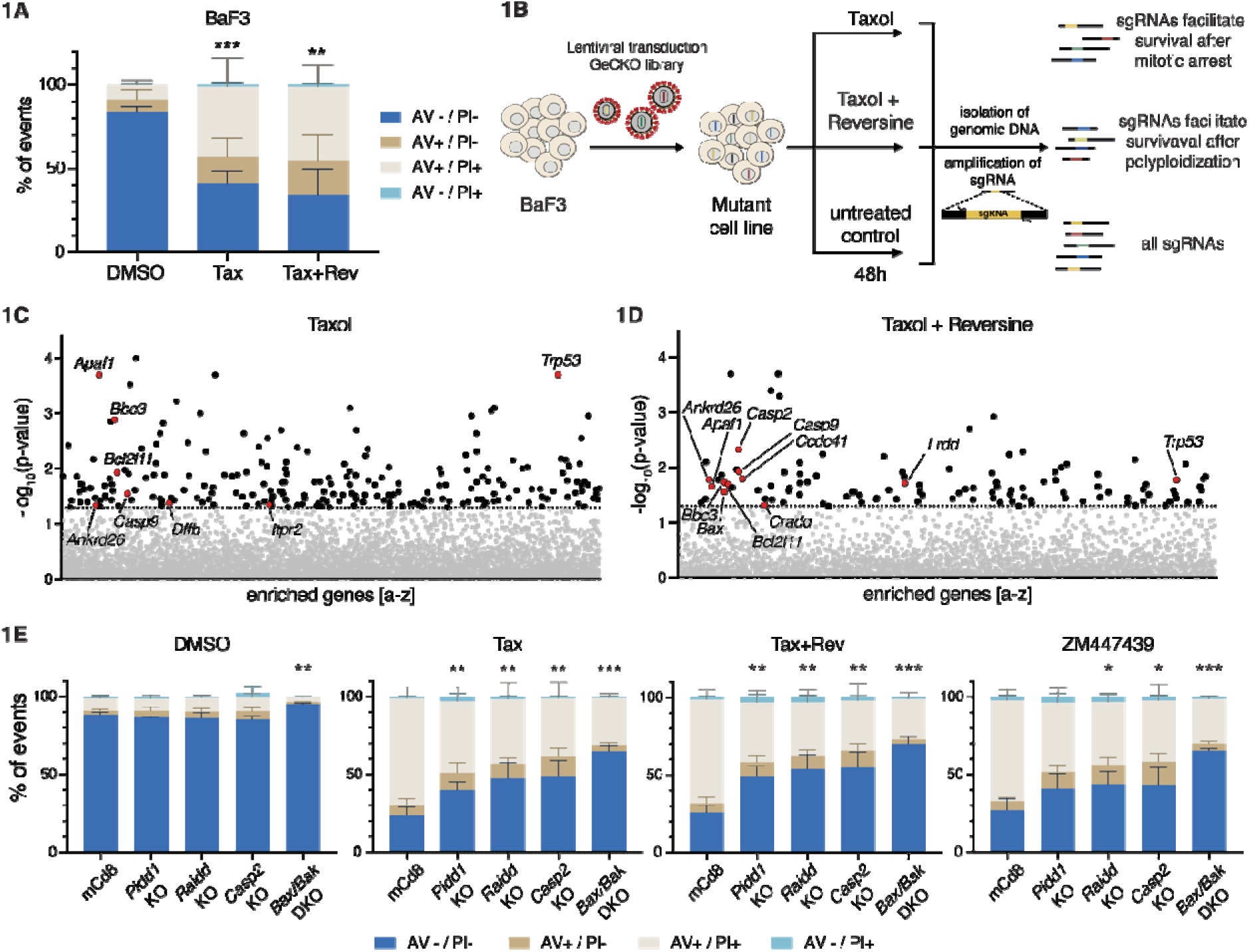
A CRISPR-Screen identifies genes involved in the induction of cell death after mitotic perturbation. **(A)** BaF3 cells were treated for 48h with 50nM Taxol (Tax) alone or in combination with 500nM Reversine (Tax+Rev) before staining with AnnexinV (AV) and Propidium Iodide (PI) followed by flow cytometric analysis. Data presented as means ± SD of the percentage of events in each staining condition. AV - / PI - = live cells; AV + / PI - = early apoptosis; AV + / PI + and AV - / PI + = late apoptosis. N = 3 independent biological replicates. Statistical significance calculated by unpaired t test on the percentage of live cells relative to the DMSO control. **(B)** Scheme of the CRISPR screen experimental setup. **(C)** Enriched sgRNAs found in surviving BaF3 cells after Taxol treatment (50nM). Horizontal dashed line indicates the significance p-value cut-off (0.05). See also Fig. S2A. **(D)** Enriched sgRNAs in surviving BaF3 cells after Taxol + Reversine treatment (50nM + 500nM). Horizontal dashed line indicates the significance p-value cut-off (0.05). See also Fig. S2B. **(E)** AnnexinV/PI staining and flow cytometric analysis of control (mCd8), Pidd1 KO, Raidd KO, Casp2 KO and Bak+Bax double KO (DKO) cells 48h after treatment with Taxol (50nM), Taxol+Reversine (50nM + 500nM), ZM447439 (2μM) or DMSO. Data presented as means ± SD of the percentage of events in each staining condition. N ≥ 3 independent biological replicates. Statistical significance calculated by unpaired t test on the percentage of live cells of each derivative clone relative to the mCd8 control for each drug treatment. See also Fig. S2C. * = p value < 0.05; ** = p value < 0.01; *** = p value < 0.001.

Consistent with these findings, BaF3 pro-B cells lacking individual PIDDosome components or the key-effectors of intrinsic mitochondrial apoptosis, BAX/BAK, were all protected from apoptosis triggered by Taxol + Reversine or, alternatively, the Aurora kinase inhibitor ZM. The same genetic knockouts (KO) also protected BaF3 cells from cell death induced by Taxol only, likely reflecting a poor propensity of these cells to maintain a strong checkpoint-dependent arrest over time (Fig. 1E and Fig. S2C).

### Caspase-2 triggers MOMP after cytokinesis failure

Our findings suggest that caspase-2 is activated in the PIDDosome to promote activation of BAX/BAK in hematopoietic cells that fail cytokinesis. For epistasis analysis and to position caspase-2 in the chain of events, we exploited a set of human Nalm6 pre-B ALL cells lacking initiator caspase-8 or caspase-9, as well as a combination of effector caspase-3 and -7 and compared their response to Nalm6 cells lacking caspase-2. Treatment with the Aurora kinase inhibitor ZM, which has been proven selective on Aurora B over Aurora A within living cells (*33*), induced apoptosis following cytokinesis failure most effectively (Fig. S1) and was used for further analysis. Cell death was assessed by Annexin V/Propidium Iodide (PI) staining and subsequent flow cytometric analysis. Notably, loss of caspase-2 provided clear protection against cell death, similar to that provided by the loss of caspase-9, or combined loss of caspase-3 and -7. In contrast, deficiency in the initiator caspase-8 failed to do so, excluding a role for death receptor-induced apoptosis in our system (Fig. 2, A and B).

**Fig. 2:**
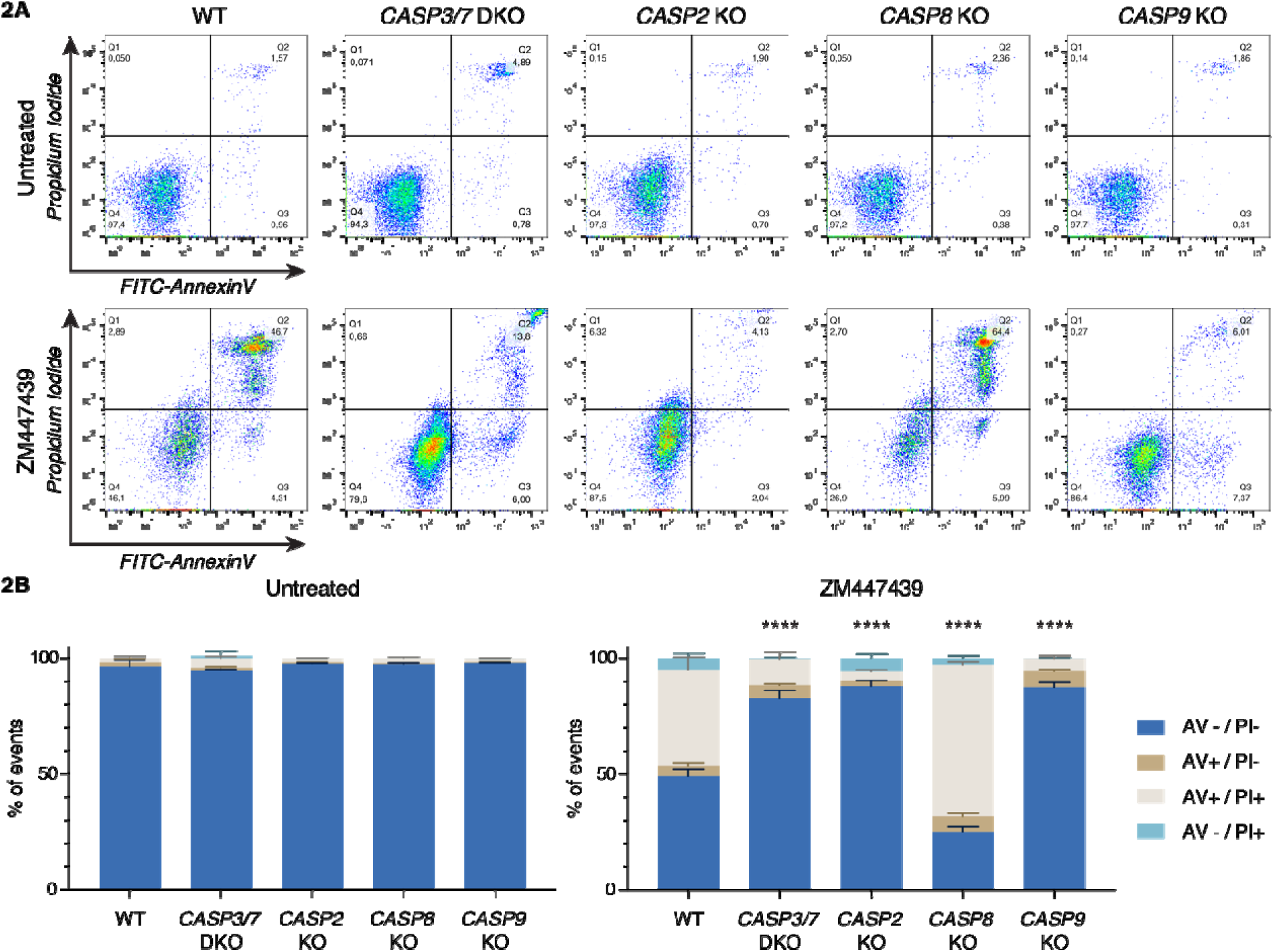
Caspase-2, caspase-9, or combined loss of effector caspase-3/-7 protects Nalm6 cells from cytokinesis failure-induced cell death, but caspase-8 does not. **(A)** Representative dot plots of AnnexinV/PI stained Nalm6 WT cells or derivative KO clones after 48h of treatment with 2μM ZM447439 (or untreated controls). **(B)** Quantification of A. Bar charts represent the means ± SD of the percentage of events in each staining condition. N ≥ 3 independent biological replicates. Statistical significance was calculated by unpaired t test on the percentage of live cells of each KO derivative clone compared to WT cells. **** = p value < 0.0001.

Based on these observations, we wondered whether caspase-2 acts upstream or downstream of caspase-9. To test this, we performed Western blotting analysis to determine caspase activation, and in particular for caspase-2 activation, we used MDM2 processing (which is known to be a substrate of caspase-2) as a readout (*34*). Given its predicted connection to caspase-2-induced cell death, we also monitored for cleavage of the BH3-only protein BID, despite not being a hit in our CRISPR screen. Western blot analysis of parental Nalm6 cells treated with ZM revealed that effector caspase-3 and -7, were processed, indicative of their activation. Moreover, we found accumulation of the processed forms of MDM2, BID, and PARP1, correlating with caspase activation and cell death initiation (Fig. 3A). However, only PARP1 cleavage was blocked in cells lacking effector caspases-3 and -7 while levels of cleaved MDM2, tBID, p53 and p21 were increased (Fig. 3A and Fig. S3). This raised the possibility that MDM2 and BID processing occurs upstream of effector caspase activation and hence by an initiator caspase (*35*, *36*).

**Fig. 3:**
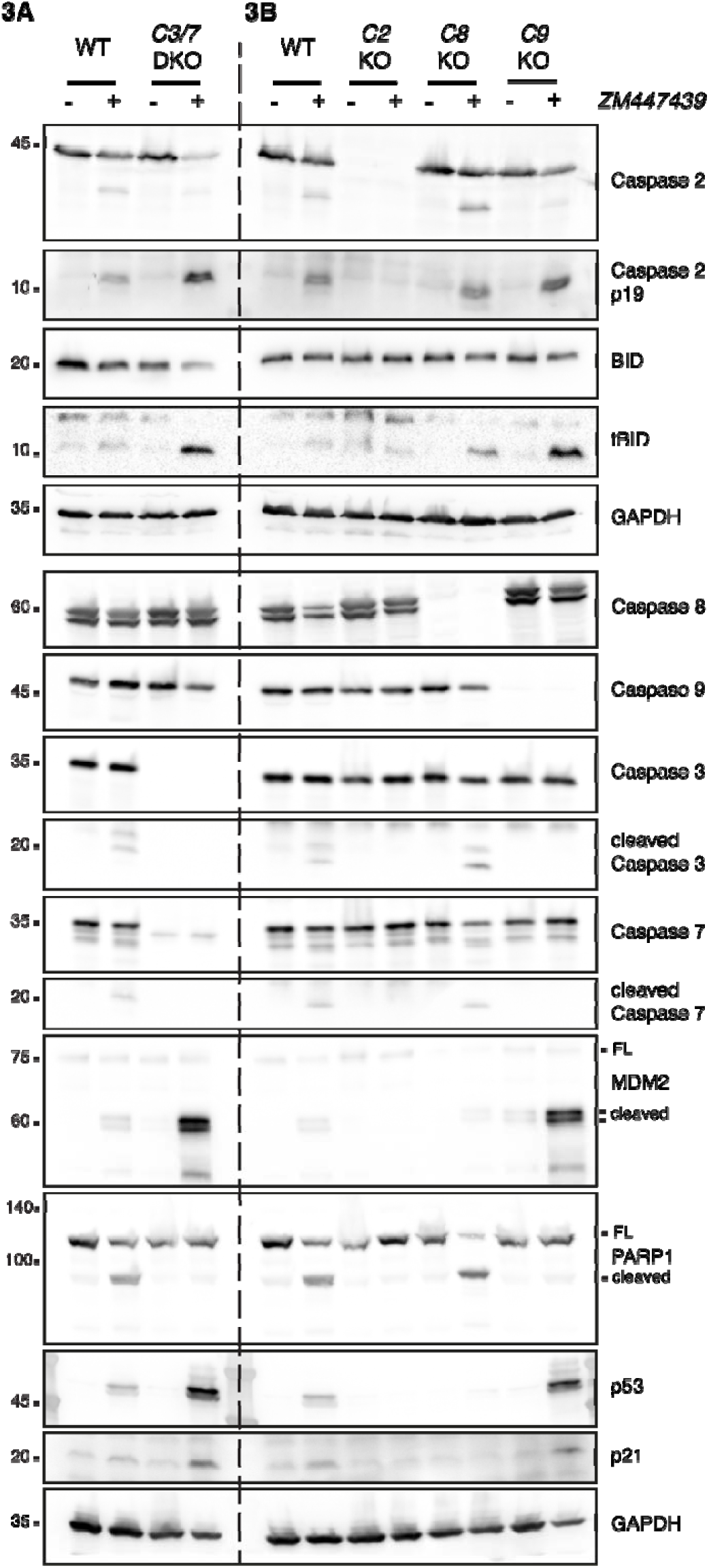
Caspase-2 acts upstream of caspase-9 to trigger apoptosis in Nalm6 cells experiencing cytokinesis failure. **(A)** Western blot analysis of Nalm6 WT and a derivative clone deficient for effector caspases-3/-7 (C3/7 DKO) after 48h of treatment with the Aurora kinase inhibitor ZM447439 (ZM, 2μM). **(B)** Western blot analysis of Nalm6 WT or knock-out clones deficient for caspase-2 (C2 KO), caspase-8 (C8 KO) or caspase-9 (C9 KO) after 48h of treatment with 2μM ZM447439 (ZM). See also Fig. S3.

To confirm if activation of caspase-2 is the cell death-initiating trigger, we next compared the impact of initiator caspase-depletion on MDM2 and BID processing. This comparison revealed that only loss of caspase-2, but not caspase-8 or -9, prevented MDM2 and tBID cleavage, despite similar cell death protection in the absence of caspase-2 or -9 (Fig. 3B). Again, impaired apoptosis in the absence of caspase-9 allowed for increased accumulation of MDM2 cleavage, p53, p21 and tBID, all of which depended on caspase-2 being present. As indicated before, caspase-8 did not contribute to any of the abovementioned phenotypes (Fig. 3B).

### Selective chemical inhibition of caspase-2 prevents substrate processing and cell death

Making use of LJ2a, a recently developed selective caspase-2 inhibitor (*37*), we compared the impact of caspase-2 or pan-caspase inhibition using Q-VD-OPh (QVD) on cell death induced by ZM or the non-genotoxic p53 activator Nutlin3 (*38*). Consistent with the specificity of LJ2a for caspase-2, Nutlin3-induced cell death was abrogated by QVD, but not LJ2a. Cell death in response to Aurora kinase inhibition was prevented by both caspase-2 and pan-caspase inhibition. In support of this view, genetic loss of caspase-9, but not caspase-2, prevented cell death induced by Nutlin3, while both mutants were protected from ZM-induced cell death. The addition of either inhibitor did not provide additional protection to caspase-2 and -9 knockout cells failing cytokinesis (Fig. 4A and Fig. S4). Western blot analysis corroborated our results using genetic caspase depletion (Fig. 3B) and revealed that cleaved MDM2 accumulated in wild type, but even more so in caspase-9-deleted cells (Fig. 4B). As expected, this was not seen in cells lacking caspase-2. Surprisingly, treatment with QVD protected against cell death in response to cytokinesis failure without preventing accumulation of the MDM2 or BID cleavage products. In contrast to this, LJ2a treatment reduced both MDM2 and BID processing in both wildtype and caspase-9 knockout Nalm6 cells alike (Fig. 4, B and C). Taken together, this indicates that caspase-2 acts upstream of mitochondria and that this activity can be selectively inhibited by LJ2a and not by QVD. The relative contribution of BID versus MDM2 cleavage to cell death execution remains instead to be established.

**Fig. 4:**
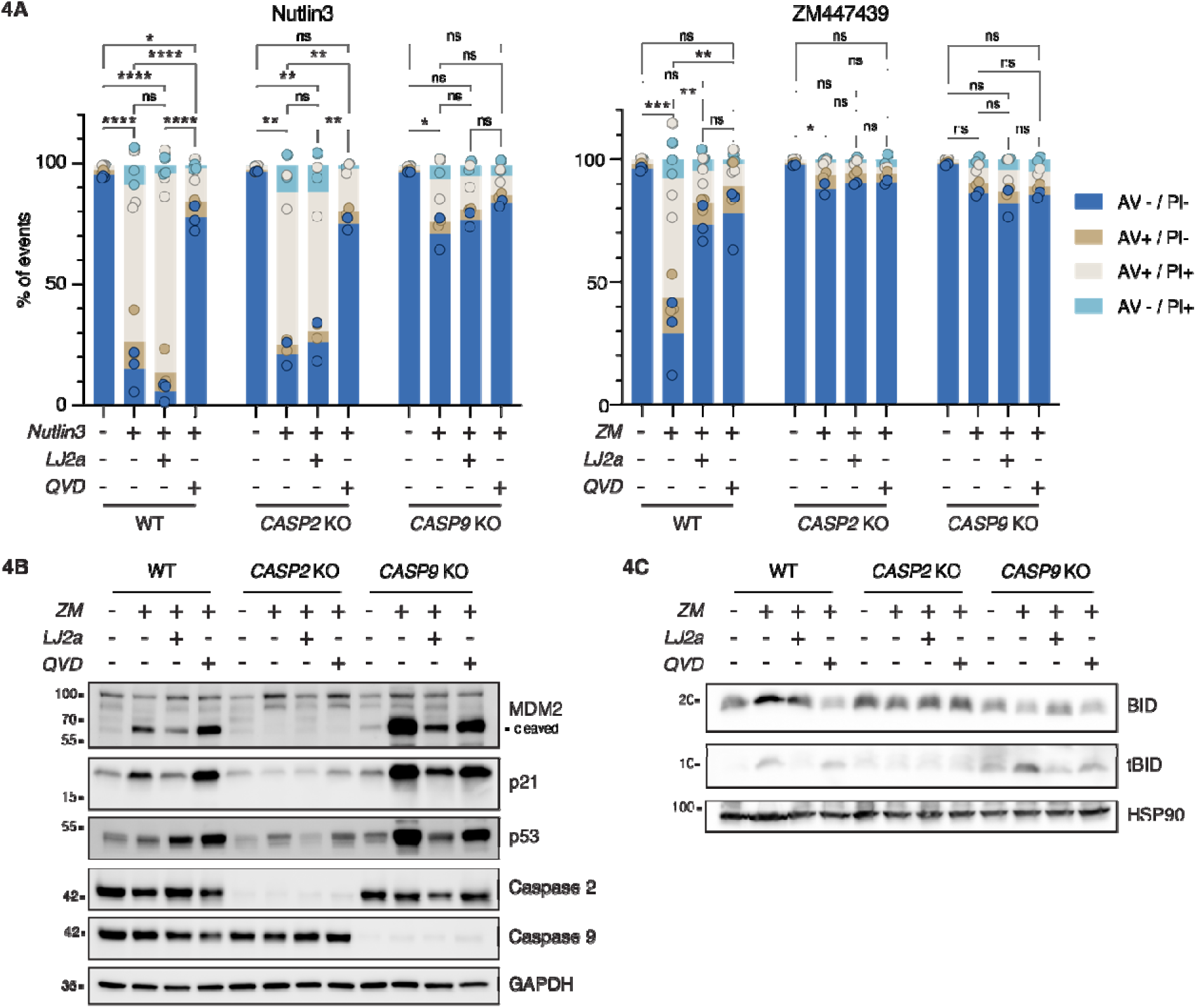
Chemical caspase-2 inhibition reduces cytokinesis failure-dependent cell death. **(A)** Nalm6 cells WT, caspase-2 KO and caspase-9 KO cells were treated with 10μM Nutlin3 or 2μM ZM447339 alone or in combination with the caspase-2 inhibitor LJ2a (10μM), the pan-caspase inhibitor Q-VD-OPh (QVD, 10μM) or left untreated for 48h before staining with AnnexinV/PI followed by flow cytometric analysis. Bar charts represent the means of th percentage of events in each staining condition, the dots represent the values for each single replicate. Statistical significance was calculated on the percentage of live cells by one-way ANOVA with Tukey’s multiple compariso correction, comparing each condition within the genotype. N ≥ 2 independent biological replicates. ns = not significant; * = p value < 0.05; ** = p value < 0.01; *** = p value < 0.001; **** = p value < 0.0001. See also Fig. S4 **(B)** Western blot analysis of Nalm6 WT, caspase-2 KO and caspase-9 KO cells after 48h of treatment with 2μM ZM447439 (ZM), alone or in combination with the caspase-2 inhibitor LJ2a (10μM) or the pan-caspase inhibitor QVD (10μM) (or untreated controls). **(C)** Western blot analysis of Nalm6 cells as described in B.

### BID is the preferred caspase-2 substrate and p53 becomes activated only when tBID production is insufficient for cell killing

Based on our results, caspase-2 acts upstream of mitochondria by stabilizing p53 via MDM2 cleavage, which may eventually induce transcription of proapoptotic effectors, such as *BBC3/PUMA*. Consistently, both *p53* and *Puma* were identified in our CRISPR screen (Fig. 1). In contrast, *Bid* was not, questioning its actual contribution to MOMP after cytokinesis failure. We therefore anticipated that p53 plays a key role in triggering cell death in response to ZM-treatment. To our surprise however, loss of p53 in Nalm6 did not protect from ZM-induced killing (Fig. 5A), prompting us to reformulate our hypothesis. Considering that both MDM2 and BID were processed in a caspase-2 dependent manner (Fig. 3B, Fig. 4, B and C and Fig. S3C), we assessed whether the two genes are able to compensate for each other. To address this possibility, we generated Nalm6 cells lacking BID and p53 alone, or in combination. Strikingly, while loss of BID alone significantly reduced ZM-induced cell death, combined loss of BID and p53 further enhanced cell death protection (Fig. 5A and Fig. S5A). This finding was confirmed in a CellTiterGlo assay, monitoring metabolic activity as a read out (Fig. 5B). Western blot analysis revealed that BID was processed into tBID following ZM treatment and that MDM2 was more processed in the absence of BID (Fig. 5C). To address whether the enhancement of MDM2 cleavage observed upon BID loss was due to delayed death or rather to enhanced capacity of caspase-2 to cleave it, we performed BID and *TP53* knockout in cell death resistant cells (i.e. devoid of caspase-9). BID loss favored MDM2 cleavage also in these experimental conditions, while loss of p53 (leading to defective MDM2 cleavage) did not alter the extent of tBID production. Taken together, our data demonstrate that the absence of BID in itself contributed to more efficient MDM2 cleavage by caspase-2 independently from longer survival, while lack of MDM2 cleavage did not alter BID cleavage propensity. Thus, BID is the preferred Capsase-2 substrate while MDM2 cleavage becomes enhanced when BID levels are reduced (Fig. S5, B and C).

**Fig. 5:**
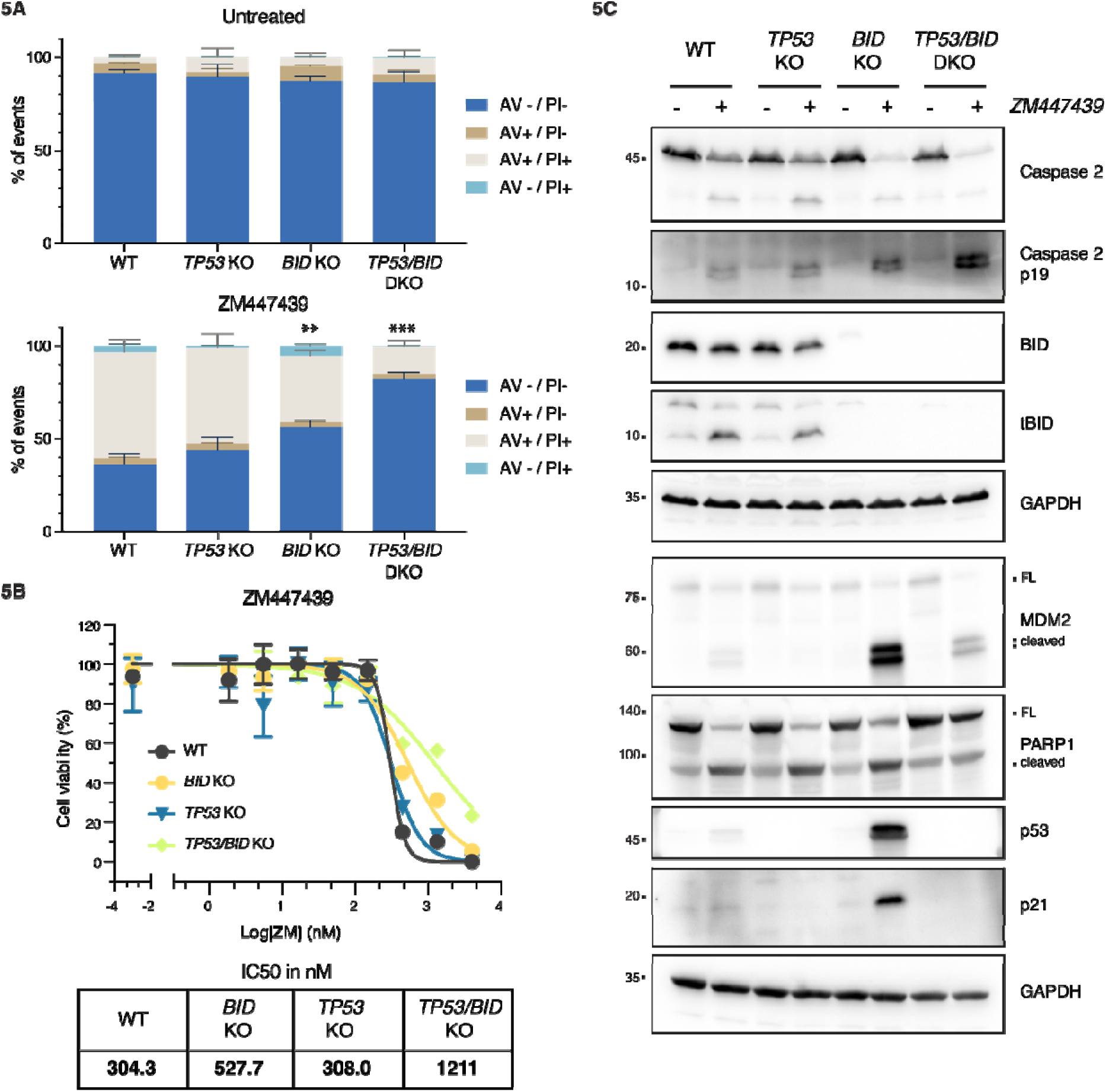
BID and MDM2 act as shared caspase-2 substrates in Nalm6 cells failing cytokinesis. **(A)** Nalm6 WT, BID KO, p53 KO or combined p53/BID (DKO) were treated for 48h with 2μM ZM447439 (or left untreated) before staining with AnnexinV/PI followed by flow cytometric analysis. Fig. S5A reports an example experiment used for quantifications shown here. Bar charts represent the means and SD of the percentage of events in each condition. Statistical significance was calculated by unpaired t test comparing each derivative clone to the WT cells. N = 3 independent biological replicates. ** = p value < 0.01; *** = p value < 0.001. **(B)** Metabolic activity of Nalm6 WT cells and derivative clones deficient for p53, BID or p53 and BID combined (DKO), as estimated by CellTiter-Glo assay after 72h treatment with the indicated concentrations of ZM447439. Lines represent the non-linear regression curves used to calculate the IC50 values reported on the bottom right table. **(C)** Western blot analysis of Nalm6 WT and derivative KO clones lacking BID, p53 or the combination after 48h of treatment with 2μM ZM447439.

### Loss of BID or caspase-9 uncovers a PIDDosome-dependent transcriptional p53 response

To better understand the interplay between BID and p53 in cell death initiation in cells failing cytokinesis, we tested different Nalm6 derivative clones lacking distinct cell death regulators after exposure to Aurora kinase inhibitor for cell death induction, ploidy levels and p53 transcriptional responses. Nutlin3 was again included as a positive control and trigger of a non-genotoxic p53 response. The absence of p53 or caspase-9 effectively protected Nalm6 cells from Nutlin3-induced cell death (Fig. 6A, Fig. S6A). As expected, Nutlin3 treatment had no effect on ploidy levels (Fig. S6A) but increased the mRNA levels of the established transcriptional p53 targets *BAX, BBC3/PUMA and p21/CDKN1A* across p53-proficient Nalm6 knockout cells (Fig. 6B). In contrast, cytokinesis failure via ZM treatment induced cell death in WT and p53 knockout cells, but was blunted in BID, caspase-2, caspase-9 knockout cells (Fig. 6C), allowing cells to become highly polyploid (Fig. S6B) without a negative impact on cell survival. Remarkably, induction of p53 targets *BBC3/PUMA* and *CDKN1A* was only detectable in cells lacking BID or caspase-9, demonstrating that this PIDDosome-dependent p53 activation is normally masked by the BID-dependent engagement of mitochondrial apoptosis (Fig. 6D).

**Fig. 6:**
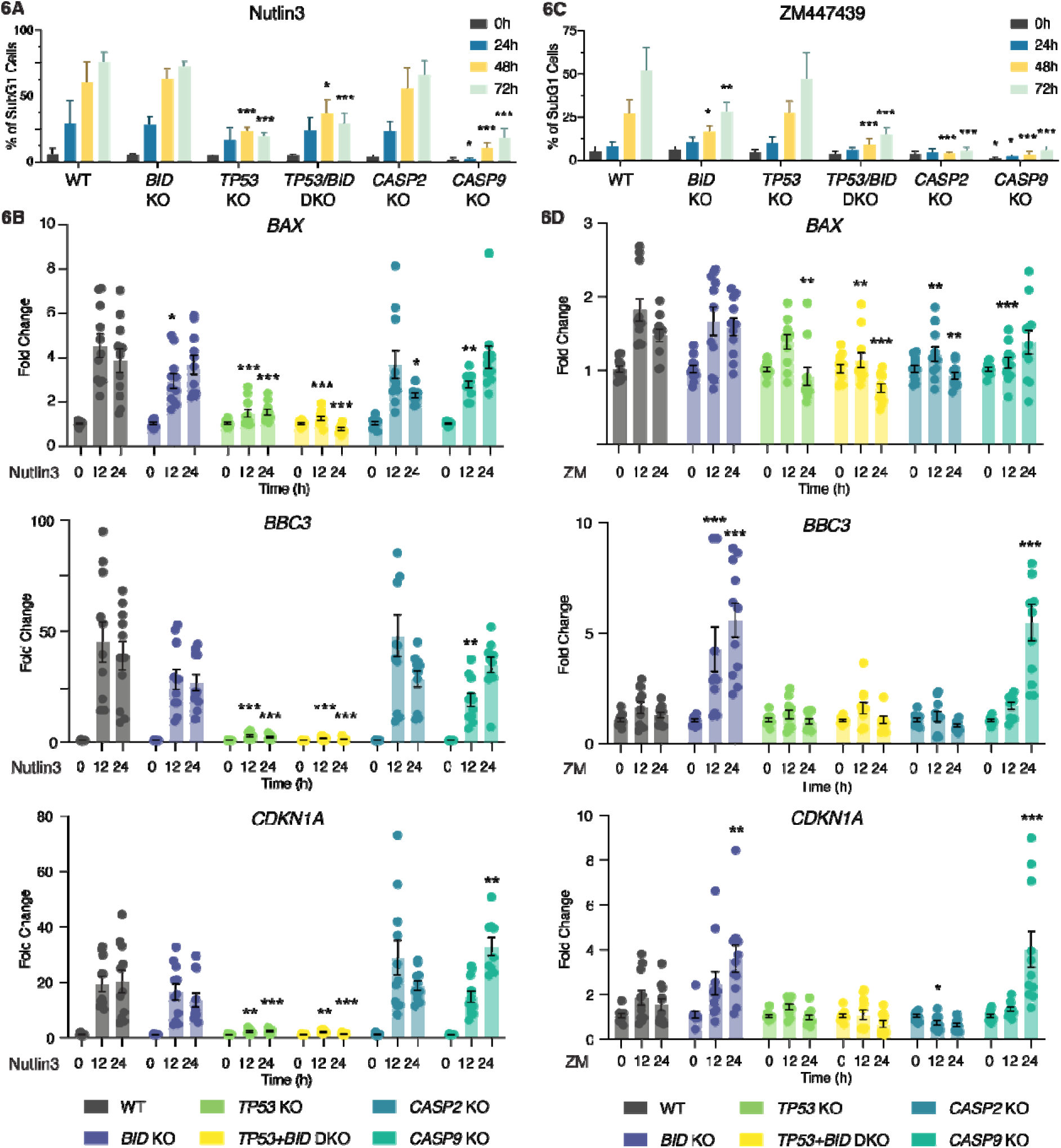
Loss of BID enables p53 dependent transcription in Nalm6 cells after cytokinesis failure. **(A)** Quantification of the percentage of subG1 cells of different Nalm6 clones at different time points after 10μM Nultin treatment. Data presented as means ± SD of N ≥ 3 independent biological replicates. Statistical significance was calculated by one-way ANOVA with Dunnett’s multiple comparison testing, comparing each time point of the KO clones to the corresponding time point of the WT sample. * = p value < 0.05; ** = p value < 0.01; *** = p value < 0.001. **(B)** RT-qPCR analysis of the p53 targets BAX, BBC3/PUMA and CDKN1A/p21 of Nalm6 WT and derivativ clones at different time points after 10μM Nutlin3 treatment. Results are normalized over the housekeeping gen GAPDH and presented as fold change over the time point 0h for each clone. Data is presented as mean ± SEM and individual points represent the values of N = 4 independent biological replicates. Statistical significance was calculated by one-way ANOVA with Dunnett’s multiple comparison test, comparing each KO clone to the WT sampl at the corresponding time point. * = p value < 0.05; ** = p value < 0.01; *** = p value < 0.001. **(C)** Same as in A, but after 2μM ZM447439 treatment. **(D)** Same as in B, but after 2μM ZM447439 treatment.

To test whether this is a more general phenomenon in hematopoietic cells, we created the same set of mutations in BL2 Burkitt lymphoma cells, which corroborated our findings in Nalm6 cells. Again, cell death induced by ZM was dependent on BID and p53 and cells became highly polyploid upon their genetic perturbation (Fig. S7, A and B). MDM2 processing, as well as p53 accumulation and p21 induction were more prominent in the absence of BID, which itself was effectively processed in wildtype and p53 mutant cells, as indicated by loss of the full-length fragment (Fig. S7D). In line with our findings in Nalm6 cells, transcriptional activation of the p53 targets, *BAX, BBC3/PUMA* and *p21/CDKN1A* was again strongest on a cell death resistant BID mutant background (Fig. S7C). Nutlin3 treatment induced the expected cell death and transcriptional response without affecting ploidy also in BL2 cells (Fig. S7, E to G).

Collectively, our data suggested that, initially, caspase-2 triggers cell death by cleaving BID into tBID to permeabilize mitochondria, yet, when this process is abrogated, a secondary transcriptional p53 response can kick-in as a fail-safe mechanism. Taken together, our findings assign caspase-2 an apical role in the mediation of p53-induced cell death, reversing the order of events in earlier models (*11*, *39*, *40*).

### Extra centrosomes are required to trigger PIDDosome-dependent apoptosis

We hypothesized that extra centrosomes were the trigger to activate PIDDosome formation and caspase-2 activation in cells that fail cytokinesis. To demonstrate this, we impaired recruitment of PIDD1 to centrosomes by generating Nalm6 cells devoid of the PIDD1 adapter at distal appendages, ANKRD26 (*29*, *30*). Immunofluorescence analysis showed that Aurora kinase inhibition led to a clear accumulation of an aberrantly high number of centrosomes in the majority of cells for all genotypes (Fig. 7, A and B). Consistently with a critical role for extra centrosomes as cell death initiators engaging PIDD1, ANKRD26 mutant cells that failed cytokinesis no longer induced caspase-2 autoprocessing, tBID formation, MDM2 processing, nor PARP1 cleavage and cell death was strongly reduced (Fig. 7, C and D), phenocopying the findings in caspase-2 mutant cells.

**Fig. 7:**
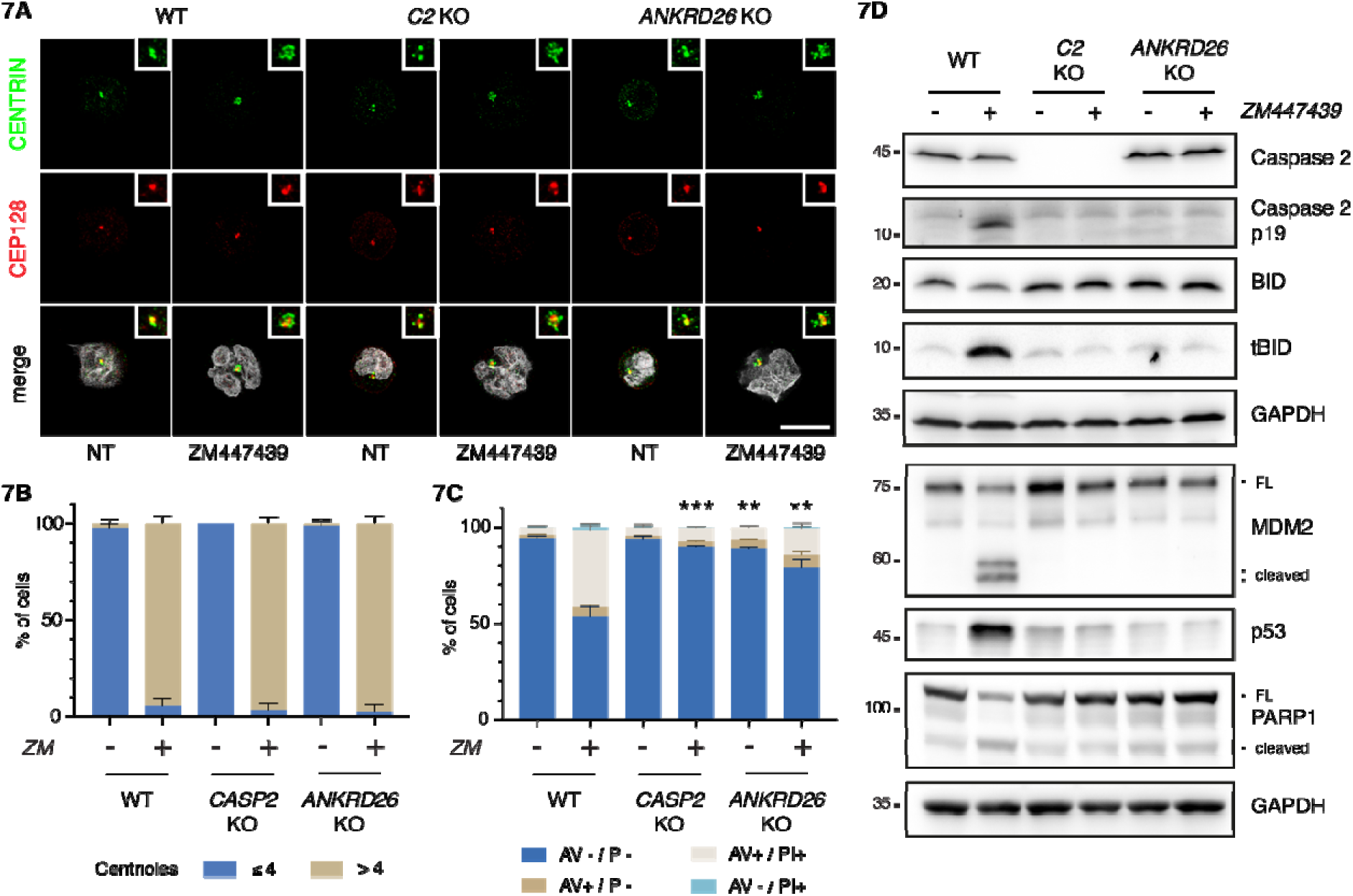
Recruitment of PIDD1 to the centrosome via ANKRD26 is necessary for cytokinesis failure-dependent cell death. **(A)** Representative immunofluorescence of Nalm6 cells WT, caspase-2 and ANKRD26 KO derivativ clones untreated or treated with 2μM ZM447439 for 48h. Cells were co-stained with the indicated antibodies: CENTRIN (in green) and CEP128 (in red). Hoechst was used to visualize the DNA (gray in the merge). Scale bar: 5µm. **(B)** Quantification of the number of centrioles of cells shown in A. Data is presented as means ± SD (i percentage) of 3 independent biological replicates. For each replicate and condition 30 cells were counted. **(C)** Percentage of Nalm6 WT, caspase-2 KO and ANKRD26 KO derivative clones undergoing apoptosis after 48 treatment with 2μM ZM447439 as detected by AnnexinV/PI staining and flow cytometric analysis. Data is presente as means ± SD of events in each staining condition (in percentage) of N = 3 independent biological replicates. Statistics were calculated by unpaired t test comparing the percentage of live cells of each knockout clone to th corresponding treatment condition in the WT sample. ** = p value < 0.01; *** = p value < 0.001. **(D)** Western blot showing Nalm6 cells WT or clones edited for caspase-2 or ANKRD26 after 48h treatment with 2μM ZM447439.

Finally, we assessed the contribution of the ANKRD26-PIDDosome axis to cell death using various drugs perturbing the fidelity of cell division. Strikingly, while a variable contribution of caspase-2 to cell death in mitosis could be appreciated upon all perturbations, the death was ANKRD26-dependent only upon perturbations that bypass the block in mitosis, promoting a direct accumulation of extra centrosomes (Fig. S8, A and B). Conceivably, all drugs perturbing mitosis promoted tBID production. Clearly, this appeared to be caspase-2-dependent only upon drug treatments promoting cytokinesis failure (namely DHCB and ZM). In contrast, tBID production in response to microtubule poisoning with Taxol and Nocodazole appeared downstream of MOMP (Fig. S8C). Together these experiments show that extra centrosomes are the trigger that can activate PIDDosome-dependent mitochondrial apoptosis in order to limit their own undesired amplification.

### BID overexpression sensitizes epithelial cells to apoptosis upon acquisition of extra centrosomes

Given that loss of BID *per se* provided a significant degree of cell death protection in response to supernumerary centrosomes in Nalm6 as well as BL2 cells, we reasoned that BID expression levels may ultimately define whether epithelial cells that frequently respond with p53-dependent cell cycle arrest may eventually commit to apoptosis induction. This prompted us to test if cell death resistant RPE1 and A549 epithelial cells can be sensitized to caspase-2 dependent cell death by overexpression of BID. Hence, we generated a doxycycline (Dox)-inducible expression system allowing for BID overexpression and treated cells with Aurora kinase inhibitor. In line with our hypothesis, upon overexpression, ZM-induced tBID production became evident in both cell lines and cells became more cell death-prone after cytokinesis failure, as indicated by increased PARP1 cleavage in Western blotting (Fig. 8, A and B) and Annexin V/PI positivity (Fig. 8, C and D and Fig. S9). Taken together, this documents a fate-switch role for BID in the cellular response to cytokinesis failure, as cell death resistant cell lines, normally responding to cytokinesis failure by displaying p53-p21-dependent cell cycle arrest, can execute some degree of apoptosis as a result of caspase-2 dependent tBID production. A graphical representation of our model is reported in Fig. 9.

**Fig. 8:**
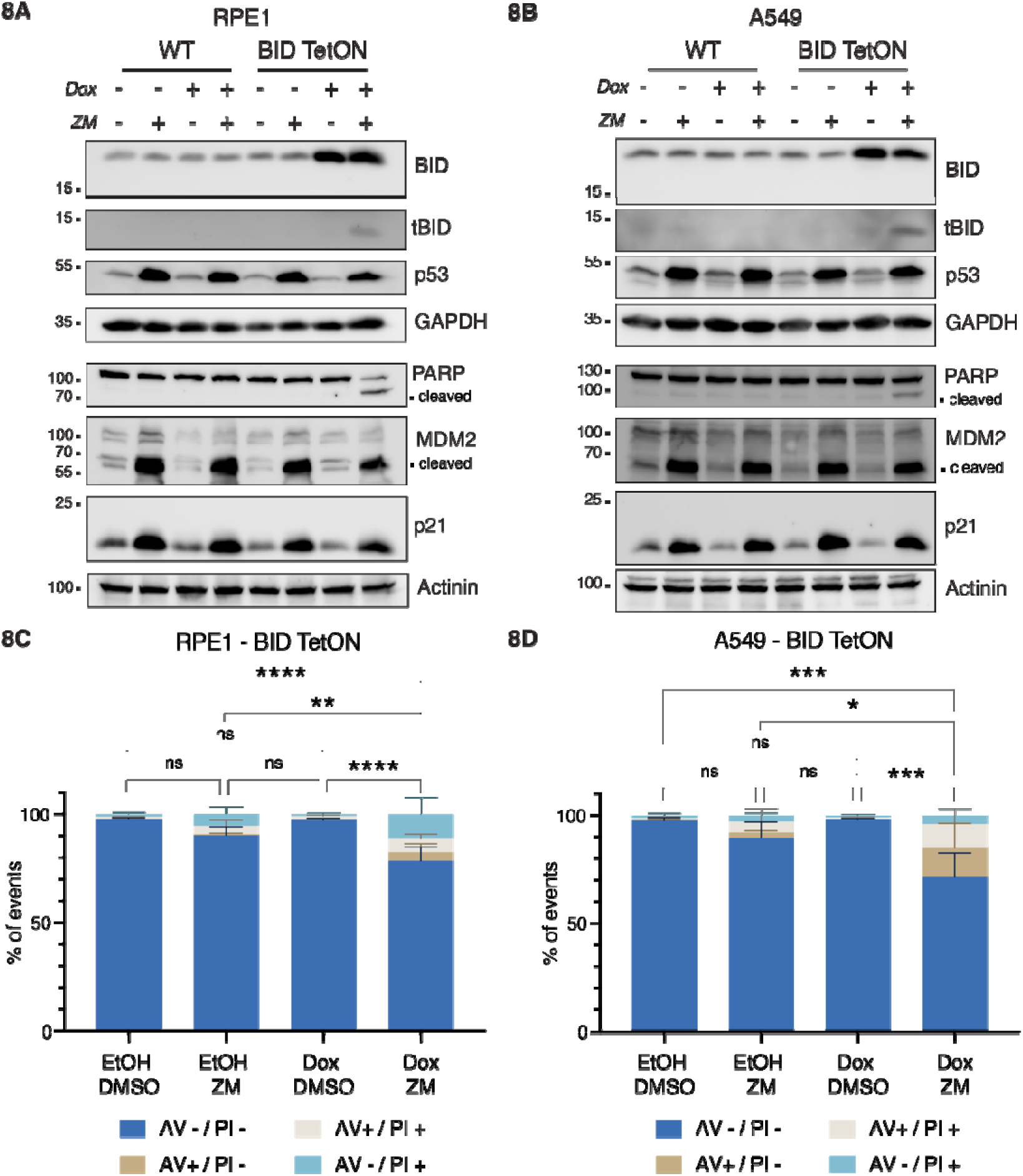
BID overexpression sensitizes epithelial cells to cell death after cytokinesis failure. **(A)** Western blot of RPE1 cells, WT or a derivative pool transduced with a doxycycline-inducible BID overexpression vector (BID TetON). Cells were treated for 24h with doxycycline (Dox; 2.5μg/ml), or solvent control (ethanol) in combination with 2μM ZM447439, or DMSO, as control. **(B)** Same as in A, but on A549 lung cancer cells. **(C)** Percentage of RPE1 BID TetON undergoing apoptosis upon treatment with 2μM ZM447439 and overexpression of BID, as detected by AnnexinV/PI staining in flow cytometric analysis. Bar charts represent the means ± SD of events in each staining condition (in percentage). Statistical significance was calculated by one-way ANOVA with Tukey’s multiple comparisons test on N = 4 independent biological replicates. * = p value < 0.05; *** = p value < 0.001; **** = p value < 0.0001; ns = not significant. **(D)** Same ad in C, but in A549 cells.

**Fig. 9:**
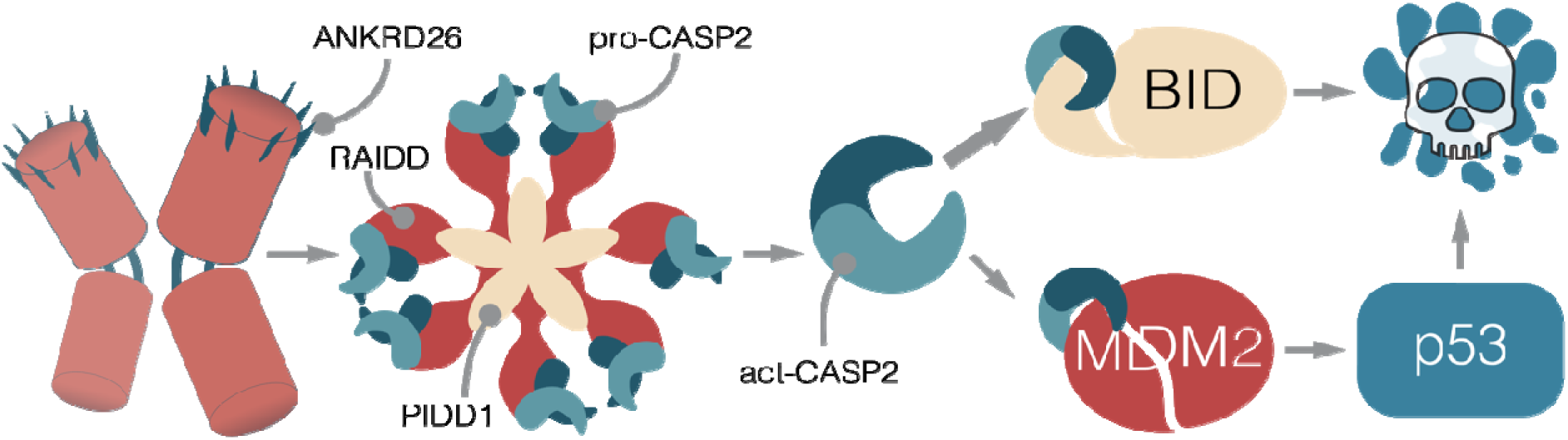
Proposed model of centrosome-dependent cell death. Hematopoietic cells failing cytokinesis acquir extra centrosomes, leading to a ANKRD26-dependent activation of the PIDDosome multiprotein complex. This leads to the autoproteolytic cleavage of caspase-2 into its active form (act-CASP2), which cleaves BID into its pro apoptotic product tBID, leading to MOMP and cell death. If BID processing is prevented, the alternative caspase-2 substrate MDM2 becomes more prominently processed, leading to p53 stabilization. The transcriptional program imposed by p53 will result in the upregulation of apoptotic effectors, ultimately leading to cell death.

## Discussion

For many years caspase-2 has been implicated in cell death initiation and execution, yet no selective trigger of its action has been identified. Several laboratories, including ours, have aimed to assess the contribution of caspase-2 in the context of apoptosis in response to DNA damage, microtubule poisons or ER stress (*16–18*). Despite having structural features of an initiator caspase, none of these studies has been able to assign an apical position to caspase-2 in any of these cell death modalities. This led to the repeated argument that it acts as a backup or amplifier in canonical, caspase-9 mediated cell death paradigms (*41*, *42*), also downstream of effector caspases (*16*). Moreover, several studies placed caspase-2 downstream of p53 (*39*, *40*, *43*). Its published role in the ill-defined process of “mitotic catastrophe” (*44*, *45*) prompted us to study its role in the context of mitotic perturbations. Building on our prior studies, identifying caspase-2 as an inducer of p53 in the context of centrosome amplification (*8*), we now provide evidence for the ability of extra centrosomes to restrict their own unscheduled duplication by the induction of PIDDosome-driven and caspase-2-dependent apoptosis as the most extreme solution to avoid consequences of potentially pathogenic polyploidy. Several lines of argument support this idea and exclude secondary effects of cytokinesis failure as drivers of cell death. First, different triggers of cytokinesis failure promote caspase-2 dependent mitochondrial apoptosis. Second, our CRISPR screen unbiasedly identified proteins involved in the formation of distal appendages that when lost provided cell death protection, including CEP83 and ANKRD26 itself, that we and the Holland lab have recently identified as the PIDD1 adapter at mature centrosomes (*29*, *30*). Finally, loss of ANKRD26 completely abrogated ZM-induced cell death in our hands. Together, this highlights the centrosome as a signalling hub for caspase activation and cell death, able to initiate pyroptosis, downstream of caspase-1 (*46*), and apoptosis as a consequence of caspase-2 activation in the PIDDosome (shown here). Notably, this pro-death role of centrosomes depends on context, e.g. bacterial infection in case of NLRP3 inflammasome activation, as well as their number and cell type, in case of centrosome amplification (discussed below).

The fact that cells acquiring extra centrosomes, for example in response to cytokinesis failure or defects in the centrosome duplication cycle, tend to lose these structures has been noted but was preferentially explained by competition of cells that either inherit a single centrosome in subsequent pseudo-bipolar cell divisions (*32*), or by the inherent anti-proliferative effects of cells carrying extra centrosomes, drastically slowing down proliferation upon p53 activation (*9*). Apoptosis as a consequence of cytokinesis defects or centrosome amplification has also been noted by some, but was never explored mechanistically and often assigned to be a consequence of p53 activation (*47*, *48*). On the other hand, the BH3-only protein BID, known for its need to be processed by different endopeptidases (most notably caspase-8) to become active, has almost exclusively been seen as a bridging element between extrinsic death receptor-induced and mitochondrial apoptosis (*27*). As such, a role for BID in other paradigms of intrinsic apoptosis was lacking until now. Our data presented here have several implications. Firstly, caspase-2 can indeed be activated upstream of mitochondria (Fig. 9), settling a longstanding debate in the cell death field (*41*, *49*). Second, caspase-2 processes BID into its active form tBID in a pathophysiological setting unrelated to death receptor signaling to drive MOMP for canonical caspase cascade activation. Also worth mentioning here is that BID has been implicated in the DNA damage response, as its loss reportedly slows down S-phase progression in response to DNA damage (*50*, *51*). This role of BID was seen as highly controversial (*25*), but may be explained by our findings that in its absence, a more effective p53 response can be mounted in blood cells that accumulate extra centrosomes, a secondary response in cells experiencing DNA damage (*5*, *52*). An interesting view supported by our work relates to the possibility that MDM2 and BID are competing substrates of caspase-2. In support of this idea, Nalm6 and BL2 BID knockouts showed a much stronger accumulation of MDM2 cleavage products in response to ZM treatment when compared to wild type cells. On the other hand, p53 functional abrogation led to blunted MDM2 cleavage, likely reflecting the inability of MDM2 gene to be transactivated by p53. Importantly, this perturbation had no impact on the propensity of the cells to cleave BID. Thus our work supports the notion that BID is the kinetically favored substrate of caspase-2, while MDM2 becomes cleaved as fail safe mechanism when tBID production is insufficient to trigger MOMP. Surprisingly, BID was not a hit of our initial genetic screen in BaF3 cells. This might simply reflect a false negative of this particular genetic screen or, as alternative, the insufficient BID expression of BaF3 cells.

Our findings also have several implications for targeted therapies currently under pre-clinical or clinical evaluation. In particular, Aurora kinase inhibitors are prominent as those with a most potent effect on cytokinesis across a wide variety of experimental conditions. It is important to note, however, that derangement of the activity of Aurora B kinase activity during cytokinetic abscission can be achieved by any trigger leading to chromosomal missegregation (*53*). Thus, the mechanism described here may be crucial for the effectiveness of a wide variety of compounds targeting mitosis, such as MPS1, CENP-E or Polo like kinase inhibitors, in addition to traditional microtubule targeting agents. Hence, assessment of caspase-2 as well as BID expression levels may help to identify cancers which respond to therapy. This concept was recently validated in non-small cell lung cancer patients with amplification of Chr22q11, which harbors the *BID* gene locus. Such cells were particularly sensitive to Aurora B or MPS1 kinase inhibition in vitro (*54*). Consistently, BID overexpression rendered A549 and RPE1 cells more susceptible to ZM treatment in our preliminary assays. In the context of cancer treatment, it appears important that loss of p53 provided only minimal protection from ZM-induced killing, raising hopes that the above-mentioned drugs may be effective in treating a broad range of tumors independent of their p53 status. This notion is backed by a report implicating induction of the BH3-only protein PUMA in response to Aurora kinase inhibition in p53 proficient as well as mutated colon cancer lines (*55*). Curiously, p53-independent *PUMA* induction has been suggested to be NF-kB dependent, but the cue driving it remained undefined. This opens the intriguing possibility that the reported ability of PIDD1 to drive NF-kB activation together with RIPK1 and NEMO in response to centrosome amplification may account for that phenomenon and contribute to cell death initiation in sensitive cell types (*56*). Moreover, the combined loss of BID and p53 provided a cell death protection comparable to loss of caspase-9 or effector caspases. This implies that p53 mutant tumors may rely heavily on BID to undergo apoptosis in the presence of extra centrosomes. Consistently, some mutations in BID that reduce its killing potency have been noted in cancer (*57*), identifying a potential drug-resistance mechanism.

Taken together, the PIDDosome emerges as a multi-pronged signaling platform that is able to instruct cells to either arrest their cell cycle via p53, release secondary messengers to alert the immune system to cells in danger of aneuploidy by engaging RIPK1, or as the most extreme measure, instruct self-destruction via BID, in settings where cell cycle arrest and senescence may not be suitable to limit cell growth and the resulting malignancy.

## Materials and Methods

### CRISPR screen

The murine precursor B cell line BaF3 expressing Cas9 were transduced with the publicly available GeCKO-V2 sgRNA library (Addgene #1000000053 (*58*)) with a multiplicity of infection (MOI) that resulted in a transduction efficacy of 10-30, thereby avoiding expression of multiple sgRNAs per cell. After 24h, transduced cells were selected with puromycin (2 μg/ml) and kept in culture for 8 days to allow the sgRNA:Cas9-mediated gene disruption to take place. Cells were treated with Taxol (50nM) and the combination Taxol (50nM) and Reversine (500nM) for 48h, while a fraction of the untreated population was snap frozen as an untreated control. Surviving cells were isolated via ficoll density gradient centrifugation (lympholyte M-cell separation Media CL5030) and snap-frozen. Generation of virus, transduction of cells, isolation of genomic DNA, PCR amplification of sgRNA sequences, and bioinformatic sgRNA enrichment analysis were performed as previously described (*59*). Raw count tables for the sgRNAs are available at (DOI: 10.5281/zenodo.13254853).

### Cell culture and drug treatments

A549, hTERT-RPE1 and HEK293T cells were cultured in DMEM (D5796 - Sigma-Aldrich) supplemented with 10% FBS (F7524 – Sigma-Aldrich), 100U Penicillin and 0.1mg/ml Streptomycin (P4333 - Sigma-Aldrich). BL2 and Ba/F3 cells were cultured in RPMI-1640 (R8758 - Sigma-Aldrich) supplemented with 10% FBS, 100U Penicillin and 0.1mg/ml Streptomycin. Ba/F3 cells were also supplemented with IL-3 derived from the filtered supernatant of WEHI-3B cells (*60*).

Nalm6 cells were maintained in RPMI medium (R8758 - Sigma-Aldrich) or Gibco (#31870-025) supplemented with 2 mM L-glutamine (Corning). In either case, media were supplemented with 10% fetal bovine serum (F7524 – Sigma-Aldrich; Gibco #1027-106), and penicillin–streptomycin solution (P4333 - Sigma-Aldrich; #15140-122-Gibco). All cells were grown at 37°C with 5% CO_2_. Cells were routinely tested for mycoplasma contamination by PCR.

Cells were treated with 10μM Nutlin-3 (10004372 – Cayman Chemical), 2μM ZM447439 (13601 - Cayman Chemical; S1103 - Selleck Chemicals), 4μM DHCB (20845 - Cayman Chemical), 100nM Nocodazole (13857 – Cayman Chemical), 50nM Taxol (10461 – Cayman Chemical), 500nM Reversine (S7588 - Selleck Chemicals), 10μM Q-VD-OPh (HY-12305 - MedChem Express; CAY15260-1 - Cayman Chemicals), 10μM LJ2a (*37*). LJ2a was purchased from Bachem Americas, Inc. (Vista, CA, USA) (custom synthesis). All drugs were resuspended in dimethyl sulfoxide (DMSO). Doxycycline (14422 – Cayman Chemical) was resuspended in ethanol and used at a final concentration of 2.5μg/ml.

### Lentiviral-based CRISPR-Cas9 gene knockout

Gene knockouts in BaF3, Nalm6 and BL2 cells were performed by lentiviral delivery of Cas9 and the sgRNA using the lentiCRISPR v2 backbone (#52961 – Addgene; a gift from Feng Zhang) (*58*). sgRNAs were cloned into the destination vector following the protocol reported on the Addgene website. The correct insertion of the sgRNA into the vector was confirmed by Sanger sequencing (Microsynth Austria). Lentiviral vectors were produced by co-transfecting the lentiCRISPR v2 containing the sgRNA with pSPAX2 and VSVg vectors in HEK293T cells using polyethylenimine (PEI - 23966-100 - Polysciences Europe GmbH) in a 1μg:2μL DNA:PEI ratio. The following day media were exchanged with fresh DMEM complete, and cells were left for 48 hours. Vectors were harvested by collecting the supernatant, removing debris by centrifugation (500g, 5min) and filtration using 0.45μm PES syringe filters.

Transduction was performed 4-24 hours after seeding the cell line of interest by adding ¼ of the media volume with the supernatant containing the viral vector. For BL2 cell transduction, protamine sulfate (1101230005 – Sigma-Aldrich) was added in combination to the viral vectors at a final concentration of 8μg/ml. Cells were left for 48 hours before starting the selection with puromycin (13884 – Cayman Chemical) at a final concentration of 1μg/ml for BaF3 cells, 1.66mg/ml for Nalm6 cells and 1.25mg/ml for BL2 cells. Media were exchanged every 48 hours, keeping the puromycin selection until the cells in the untransduced well were all dead. Polyclonal pools were expanded, and the gene knockout efficiency was tested by Western blotting. To select cells with the highest editing efficiency on BL2 cells, polyclonal pools were seeded on a 96 well format to a concentration of ∼ 5 cells/well and allowed to grow. After expansion, the sub-pools were tested for knockout efficiency via western blot and only the sub-pools showing the highest reduction on the protein levels were kept for further experiments. For the pools transduced with constructs targeting TP53, a further selection using 10μM Nutlin3 for 48h was performed to select only for those cells harboring a p53 deletion or mutation.

### RNP-based CRISPR-Cas9 gene knockout

Nalm6-KO cell lines were generated using an RNP-based CRISPR/Cas9 approach, as outlined by Ghetti et al (*61*) (without the incorporation of a single-strand DNA homology template and NU-7741 treatment). Isogenic clones for all cell lines were obtained through limiting dilution. The presence of gene-disrupting INDELs in edited cells was validated by Sanger sequencing of PCR products encompassing the crRNA recognition site, followed by analysis using the Inference of CRISPR Edits (ICE) tool (https://ice.synthego.com) (*62*). Caspase-8, -9, Caspase 3,7 (DKO) and −3,6,7 (TKO) cell lines were described before (*63*). Caspase-9 p53 KO cells were electroporated as described before. After 2 days, p53 KO cells were enriched by Nutlin-3a treatment (10 μM for 1 week). p53 Knockout efficiency was verified using the Inference of CRISPR Edits (ICE) tool. In this case, single cell clones were obtained through limiting dilution.

### Cell cycle profiling – SubG1 analysis

Cells were collected at the desired time point after the treatment and washed once with PBS before fixation in 70% cold ethanol and stored for at least 4 hours at -20°C. Subsequently, fixed cells were pelleted (1000g, 3min) and washed twice in PBS before resuspension in DNA staining solution, composed by 10μg/ml propidium iodide (PI – 14289 – Cayman Chemical), 100μg/ml RNase A (10109169001 - Roche) in PBS. The volume of DNA staining solution varied depending on the size of the cell pellet, to not exceed a concentration of 1×10^6 cells/ml. Samples were filtered using 50μm Filcon (340631 – BD Biosciences) and acquired using a LSR Fortessa Cell Analyzer (BD Biosciences). Analysis was performed using FlowJo v10, setting the gates to remove doublets before determining the threshold for the subG1 population.

### Cell viability assay

Nalm-6 cells were seeded in 96-well plates at a density of 5000 cells per well, incubated for 24 hours and subsequently treated with DMSO or ZM447439. The experiments were performed in biological triplicates and ZM447439 was serially diluted to 4000, 1333, 444.4, 148.1, 49.39, 16.46 nM before cell treatment. Cells were then incubated for 72h and the ATP content of every well was assessed on the Plate Reader Platform Victor X3 model 2030 (Perkin Elmer) using the CellTiter-Glo assay (Promega G7573), following the manufacturer’s protocol. Data was analyzed using GraphPad Prism v10. All data points were normalized to every cell line’s respective mean luminescence DMSO control. Dose-response curves were generated by non-linear regression curve fitting from which then IC50 values were derived.

### RNA extraction, cDNA conversion and RT-qPCR

Cells were collected after treatment and washed once with PBS before snap freezing. Pellets were resuspended in 100μl PBS before adding an equal volume of TRIzol reagent (15596026 - Invitrogen). After 5 minutes of incubation, chloroform was added (1/5 of TRIzol volume), samples were vortexed for a few seconds and incubated for 3 minutes before centrifugation (12000g, 4°C, 15 min). The clear, upper phase was collected and transferred to a new tube, added with 0.5ml of isopropanol and 1μl of GlycoBlue Coprecipitant (AM9516 - Invitrogen) before the next centrifugation step (12000g, 4°C, 10 min). The supernatant was removed and the RNA pellet was washed once with 75% ethanol. After another centrifugation step (12000g, 4°C, 10 min) ethanol was removed and the pellet was left to dry for 5 minutes before resuspension in molecular biology grade water (46-000-CV - Corning). RNA concentration was determined using a Nanodrop 2000c (Thermo Scientific). Depending on the sample concentration, 750 to 1500ng of RNA were used for retrotranscription to cDNA. cDNA conversion was performed using the RevertAid First Strand cDNA Synthesis Kit (K1622 – Thermo Scientific) according to the manufacturer’s instructions. Samples were diluted with molecular biology grade water to a final concentration of 10ng/μl, assuming 100% retrotranscription efficiency.

RT-qPCR was performed starting from 20ng of cDNA for each target gene and in technical duplicate or triplicate using the qPCRBIO SyGreen Blue Mix (PB20.17-20 – PCR Biosystems) following the manufacturer’s instruction. Amplification and detection were performed using CFX Opus 384 Dx (Bio-Rad) and Ct values were determined using CFX Maestro 2.3 (Bio-Rad). Fold changes were calculated over the housekeeping gene and the corresponding control treatment using the 2^-ddCt method using Microsoft Excel. Plots were produced using Prism v10 (GraphPad).

### Western Blot

After treatment with the desired drug (or DMSO control), cells were collected, washed once with PBS and pellets were snap frozen. An appropriate volume of Lysis Buffer (50mM TrisHCl pH 7.4, 150 mM NaCl, 0.5 mM NP40, 50mM NaF, 1mM Na_3_VO_4_, 1mM PMSF, 30μg/ml DNaseI (DN25-10MG – Sigma Aldrich), 1 tablet/10ml of cOmplete protease inhibitor, EDTA-free (4693132001 - Roche)) was added to the cell pellet for 40 minutes before centrifugation (16000g, 4°C, 12min). Superantants were collected and quantified using Pierce BCA Protein assay (23227 – Thermo Scientific). Depending on the sample quantification and the proteins to be detected, a volume corresponding to 20 to 60μg of proteins was used for SDS-PAGE using Tris/Glycine/SDS buffer. Blotting was performed in wet conditions using a Tris/Glycine buffer added with 20% ethanol on nitrocellulose membranes (GE10600002 - Cytiva). PVDF (GE10600029 – Cytiva) membranes were used for the detection of tBID. Blocking was performed in 5% milk (T145.3 – Carl Roth) in TBS-T. Antibodies were all diluted in 1% milk in TBS-T; primary antibodies were all incubated overnight at 4°C whereas secondary antibodies were incubated for 1 hour at RT. Detection was performed by incubating the membranes with ECL Select Western Blotting Detection Reagent (GERPN2235 – Cytiva) for 3 minutes before detection using a ChemidocMP (Bio-Rad) or Alliance LD2 Imaging System (UviTec Cambridge).

### Immunofluorescence

Cells, treated or untreated, were counted and 100.000 cells were plated on 12 mm glass coverslips pre-coated with 100 μg/mL Poly-D-lysine. Then cells were left 30 minutes at RT or 4°C and fixed and permeabilized with absolute ice-cold methanol for at least 20 min at −20°C.

Cells were rinsed with PBS, blocked with 3% w/v BSA in PBS for 20 min and stained for 1 h at room temperature with primary antibodies diluted in blocking solution. Cells were washed with PBS and incubated with fluorescent secondary antibodies for 45 min at room temperature. DNA was stained with 1μg/ml Hoechst 33342 (Invitrogen). After incubation, cells were rinsed with PBS and ddH2O, and mounted using ProLong Gold Antifade Reagent (Invitrogen).

Images for quantification were acquired using a Nikon Eclipse Ti2 inverted microscope, equipped with a CrestOptics X-Light V2 spinning disc module, a Lumencor SpectraX light engine and an Andor iXon Ultra 888 EMCCD camera using a plan apochromatic 100×/1.45 oil immersion objective. Representative images were acquired on a Leica TCS SP8 microscope using a 63×/1.4 oil objective with Lightening mode (adaptive as “Strategy” and ProLong Gold as “Mounting medium”) to generate deconvolved images. Images were processed using Fiji and displayed as maximum intensity projections of deconvolved z-stacks.

### AnnexinV-PI staining

Cells were collected after the treatments and washed twice with PBS before resuspension in 100μl of Annexin Binding Buffer (140mM NaCl, 2.5mM CaCl_2_, 10mM Hepes pH 7.4). Samples were added with 5μl of FITC AnnexinV antibody (640906 – Biolegend) and 10μl of PI (0.5mg/ml). Alternatively, samples were stained using the FITC AnnexinV Apoptosis Detection Kit I (BD Pharmingen, 556547) following the manufacturer’s instructions. After vortexing, samples were incubated for 15 minutes at room temperature in the dark before adding 400μl of Annexin Binding Buffer and proceeding with the acquisition using a LSR Fortessa Cell Analyzer (BD Biosciences) or a Symphony A1 cytometer (BD Biosciences). Data were processed using FlowJo v10 (FlowJo, LLC) and summary plots were obtained using Prism 10 (GraphPad).

### BID overexpression

BID sequence was amplified from hTERT-RPE1 cells cDNA using primers containing the restriction sites for NheI and AgeI. The PCR fragment, as well as the destination vector pCW57.1 (Addgene plasmid # 41393, a gift from David Root) were digested using NheI-HF (R3131S - NEB) and AgeI-HF (R3552S - NEB). The PCR fragment was subsequently purified using the Wizard SV Gel and PCR Clean-Up System (A9281 - Promega) according to the manufacturer’s instructions. The vector was dephosphorylated using the rSAP (M0371S - NEB), loaded on a gel and the correct band was purified using Wizard SV Gel and PCR Clean-Up System. Ligation of the PCR product with the purified destination vector was performed using T4 DNA Ligase (M0202S - NEB) and transformed in Stbl3 bacteria. Minipreps from selected colonies were sequenced by Sanger sequencing (Microsynth Austria) to ensure the correct cloning of the insert.

Viral vectors and transduction were performed as described above (see the Lentiviral based CRISPR-Cas9 gene knockout section), with the only exception of plasmid pMD2.G (Addgene plasmid # 12259, a gift from Didier Trono) being used instead of VSVg. A549 cells were selected with 2.5mg/ml puromycin and hTERT-RPE1 with 10mg/ml puromycin and kept in selection until control cells were all dead. After expansion, the pools of transduced cells were co-treated with 2.5μg/ml doxycycline (Cay14422-1 – Cayman Chemical) or the corresponding volume of ethanol and ZM447439 for 24 hours before collection and prepared for Western blot analysis. For AnnexinV/PI staining, A549 and RPE1 cells were co-treated with doxycycline (or ethanol) and ZM447439 (or DMSO) 24h after seeding.

### Statistical Analysis

Data are presented as means and standard deviation or standard error of the mean as reported in the legends for each panel. All the statistical analyses were performed using GraphPad Prism v10 (GraphPad Software). Details on the statistical tests, number of replicates and significance intervals are reported in the figure legends.

---

**Fig. S1:**
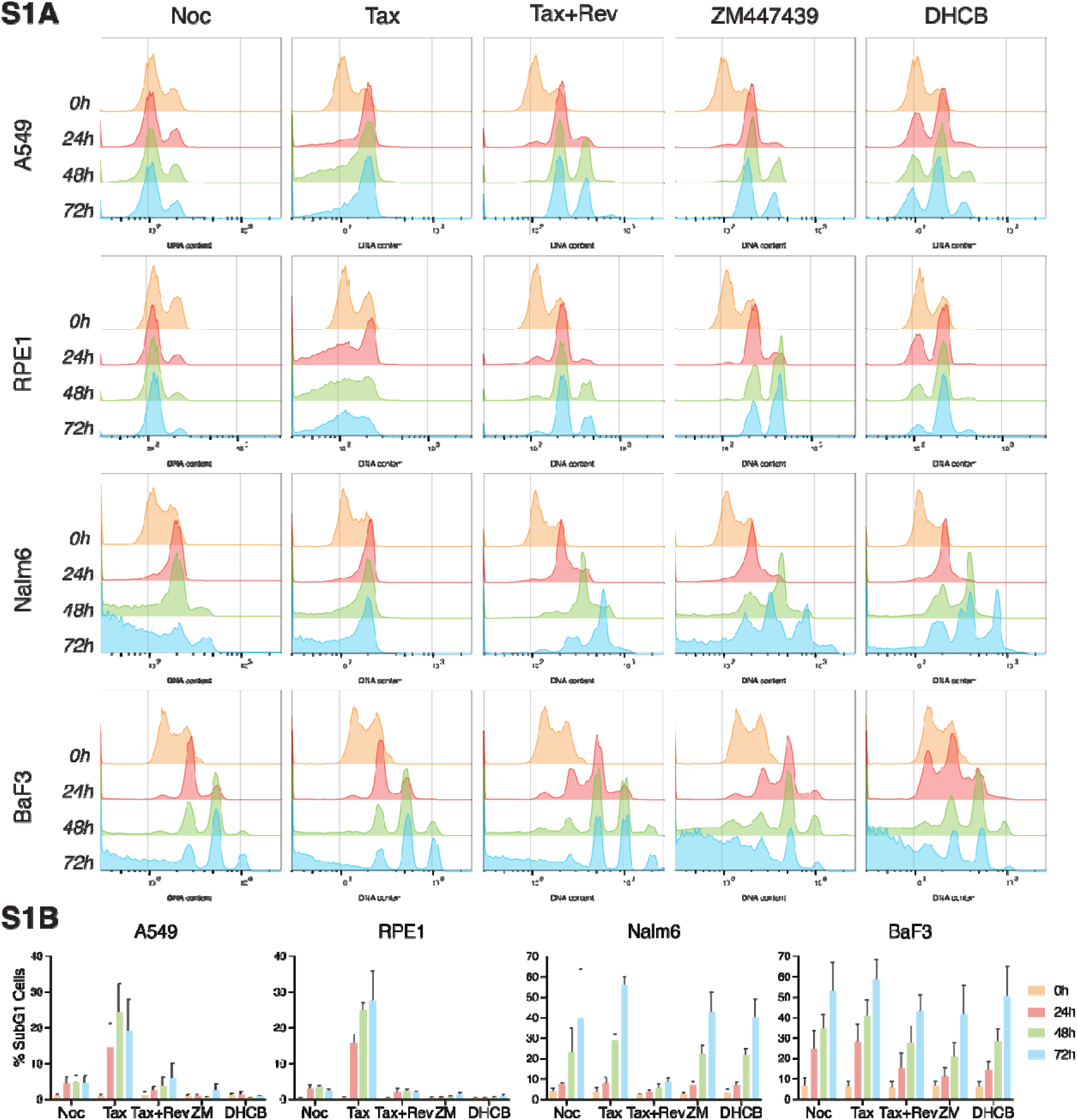
The outcome of mitotic errors is cell line dependent. **(A)** Representative DNA content profiles of A549, RPE1, Nalm6 and BaF3 cell lines exposed to different drugs interfering with mitotic progression (100nM Nocodazole, Noc; 50nM Taxol, Tax) and cytokinesis (50nM Taxol + 500nM Reversine, Tax+Rev; 2μM ZM447439, 4μM DHCB) as detected by propidium iodide staining and flow cytometric analysis. For each treatment, three different time points were analyzed. **(B)** Quantification of the subG1 events from A. Data is presented as mean ± SD of N ≥ 3 independent biological replicates for each condition.

**Fig. S2:**
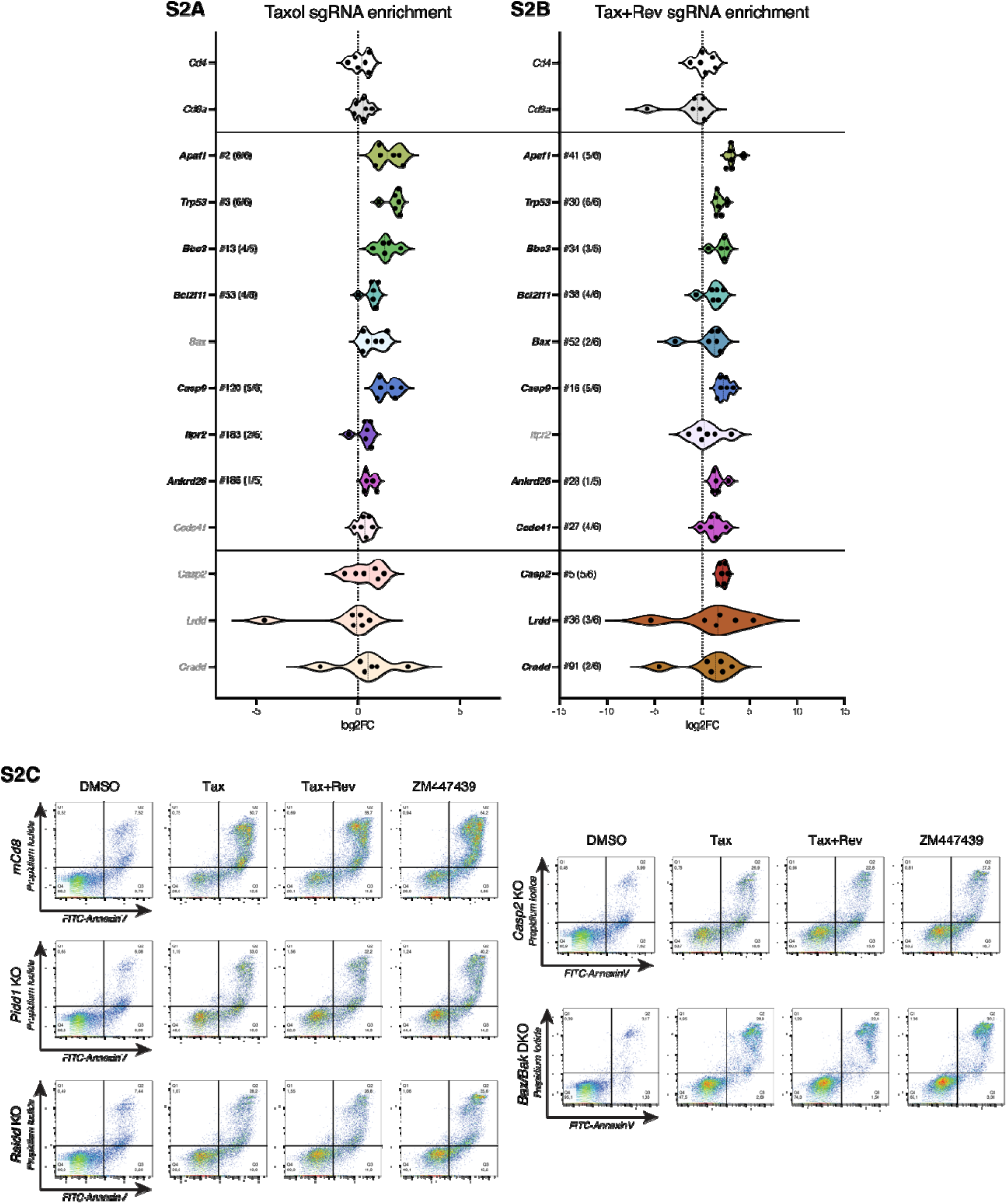
BaF3 CRISPR Screen sgRNA distribution for selected genes and validation of the central role of the PIDDosome in mitotic error-induced apoptosis. **(A)** log2 fold change (log2FC) distribution of significantly enriched sgRNA targeting Trp53 and other genes involved in apoptosis after Taxol treatment (50nM). Each dot represents sgRNA. Cd4 and Cd8a were included as negative controls. Genes marked in bold are significantly enriched (p value < 0.05). The number following the # indicates enrichment ranking of the gene whereas the values in brackets indicate the number of enriched sgRNAs targeting that gene. **(B)** log2 fold change distribution of significantly enriched sgRNA targeting the PIDDosome components, Trp53 and genes involved in apoptosis after Taxol+Reversin treatment (50nm + 500nM). Each dot represents a sgRNA. Cd4 and Cd8a were included as negative controls. Genes reported in bold are significantly enriched (p value < 0.05). The number following the # indicates gene enrichment ranking whereas the values in brackets indicate the number of enriched sgRNAs targeting that gene. **(C)** Representative dot plot examples of flow cytometric AnnexinV/PI analyses of BaF3 cells lacking Pidd1, Raidd, Casp2, Bak and Bax, or harboring a control guide RNA targeting mouse Cd8 (mCd8). Cells were treated for 48 hours with 50nM Taxol, 50nM Taxol + 500nM Reversine and 2μM ZM447439. Quantification is shown in Fig.1E.

**Fig. S3:**
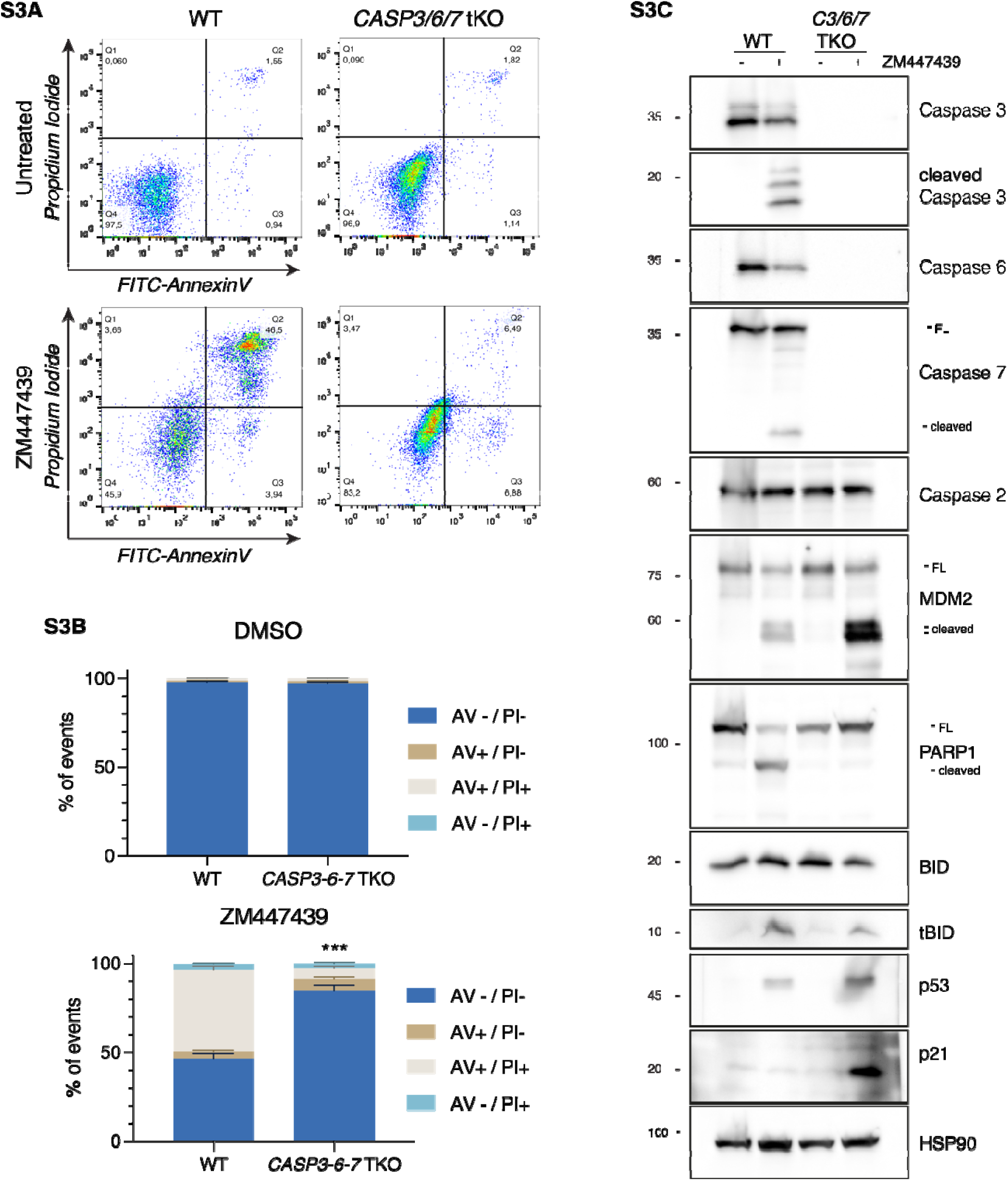
Caspase-3,-6,-7 triple KO cells show enhanced survival and increased MDM2 processing after ZM447438 treatment. **(A)** Representative dot-plots of AnnexinV/PI stained Nalm6 WT cells and a derivative clone lacking effector caspases-3,-6,-7 after 48h of treatment with 2uM ZM447439 (or untreated controls). **(B)** Quantification of A. Bar charts represent the means ± SD of the percentage of events in each staining condition. N = 3 independent biological replicates. Statistical significance was calculated by unpaired t test on the percentage of liv cells of the caspase-3,-6,-7 triple KO clone compared to WT cells. *** = p value < 0.001 **(C)** Western blot analysis of WT and effector caspases-3,-6,-7 TKO cells after 48h of treatment with the Aurora kinase inhibitor ZM447439 (2μM).

**Fig. S4:**
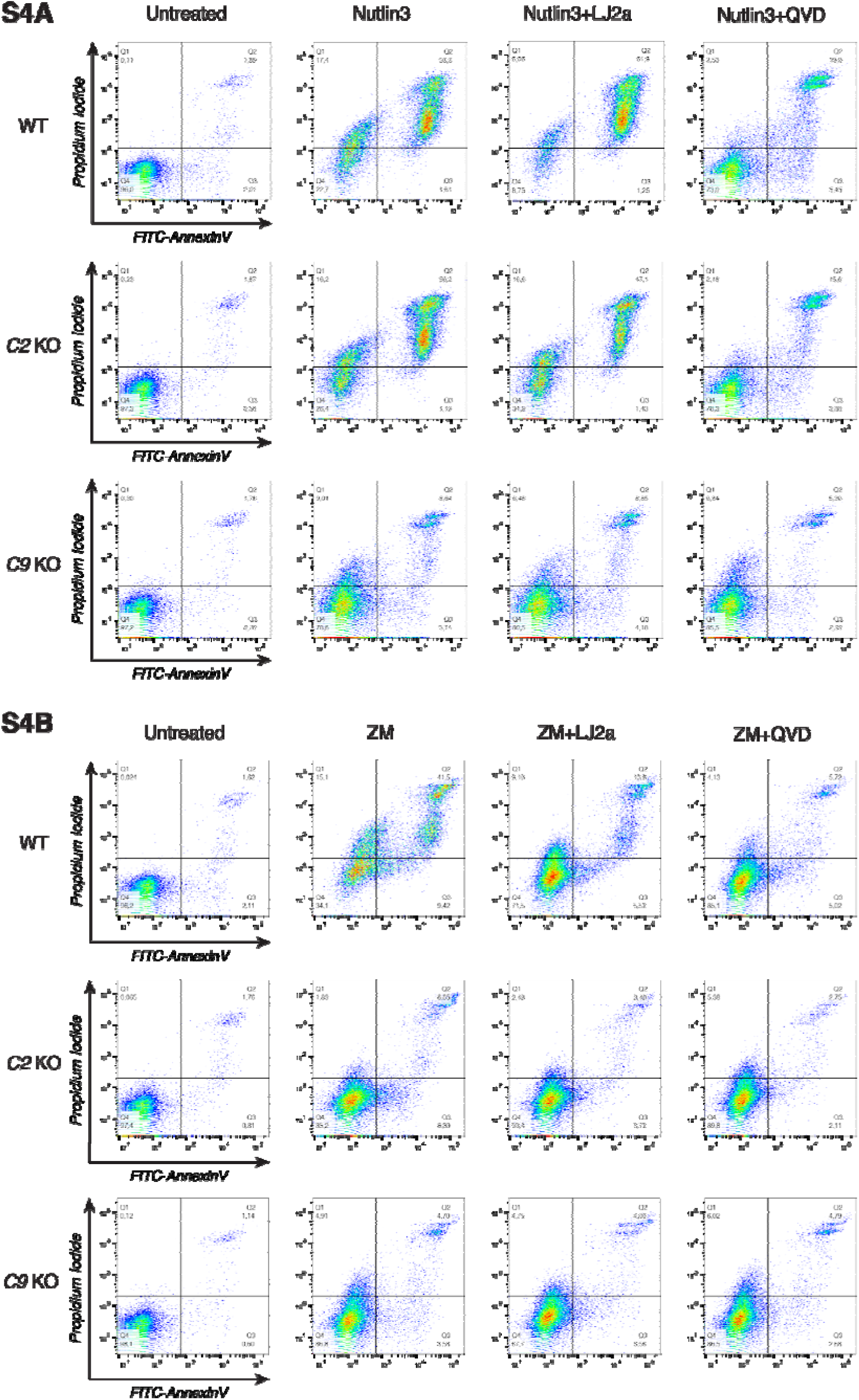
Loss or inhibition of caspase-2 prevents only cytokinesis failure-dependent apoptosis but not Nutlin3-induced cell death. Representative dot plots examples of AnnexinV/PI staining of Nalm6 WT cells or derivative clones edited for caspase-2 or caspase-9 after 48 hours of treatment with 10μM Nutlin3 (panel A) or 2μM ZM447439 (panel B) alone or in combination with the caspase-2 inhibitor LJ2a (10μM) or the pan-caspase inhibitor QVD (10μM). Quantification is shown in Fig. 4A.

**Fig. S5:**
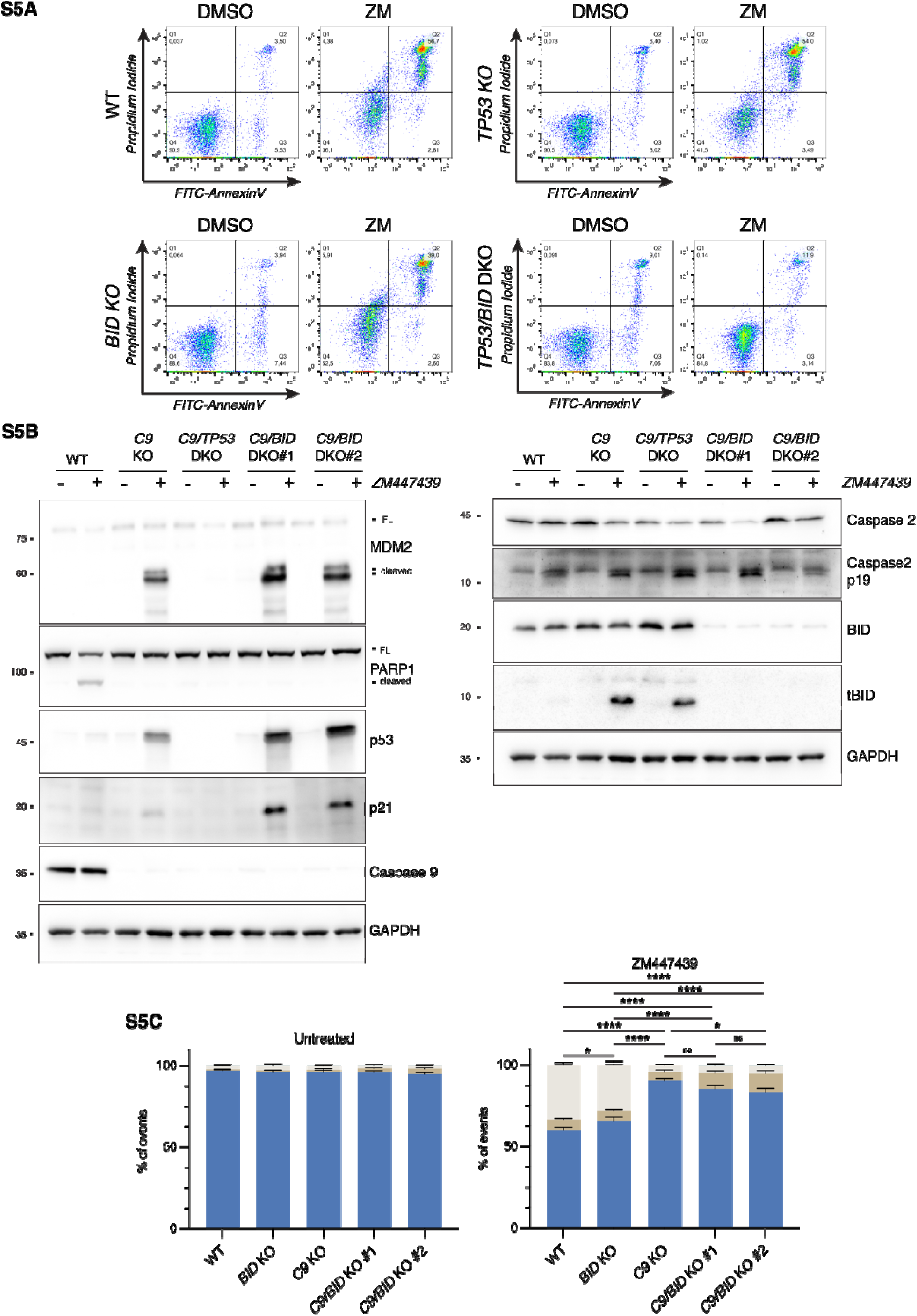
BID is the kinetically favored caspase-2 substrate. **(A)** Representative dot-plots of AnnexinV/PI staining of Nalm6 WT cells or derivative clones lacking p53, BID or both after 48 hours of treatment with 2μM ZM447439. Quantification is shown in Fig. 5A. **(B)** Western blot of Nalm6 WT and derivative clones lacking caspase-9 only, caspase-9 in combination with p53 (C9/TP53 DKO) or BID (C9/BID DKO, two independent clones #1 and #2) after 48h of treatment with ZM447439 (ZM, 2μM). **(C)** Quantification of AnnexinV/PI staining and flow cytometric analysis of Nalm6 WT and derivative clones described in B treated for 48h with 2μM ZM447439. Data are presented as means ± SD of the percentage of events in each condition. N = 3 independent biological replicates. Statistical significance was calculated by one-way ANOVA with Tukey’s multiple comparison test on the percentage of live cells within each genotype. ns = not significant; * = p value < 0.05; **** = p value < 0.0001.

**Fig. S6:**
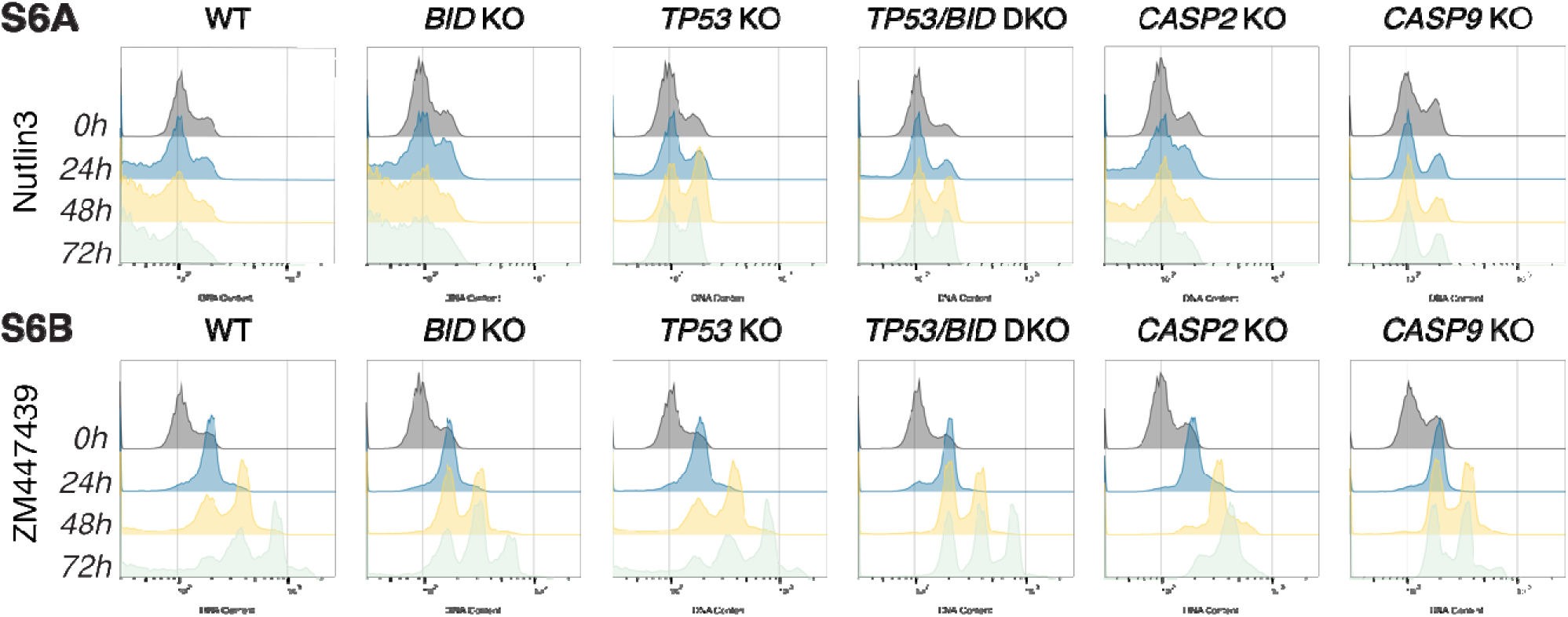
Combined loss of BID and p53 impairs cytokinesis failure-induced cell death. **(A)** Representative DNA content profiles of Nalm6 WT cells and derivative clones at different time points after 10μM Nutlin3 treatment. Quantification of the subG1 events shown in Fig. 6A. **(B)** Same as in A, after cytokinesis failure induced by 2μM ZM447439. Quantification of the subG1 events shown in Fig. 6C.

**Fig. S7:**
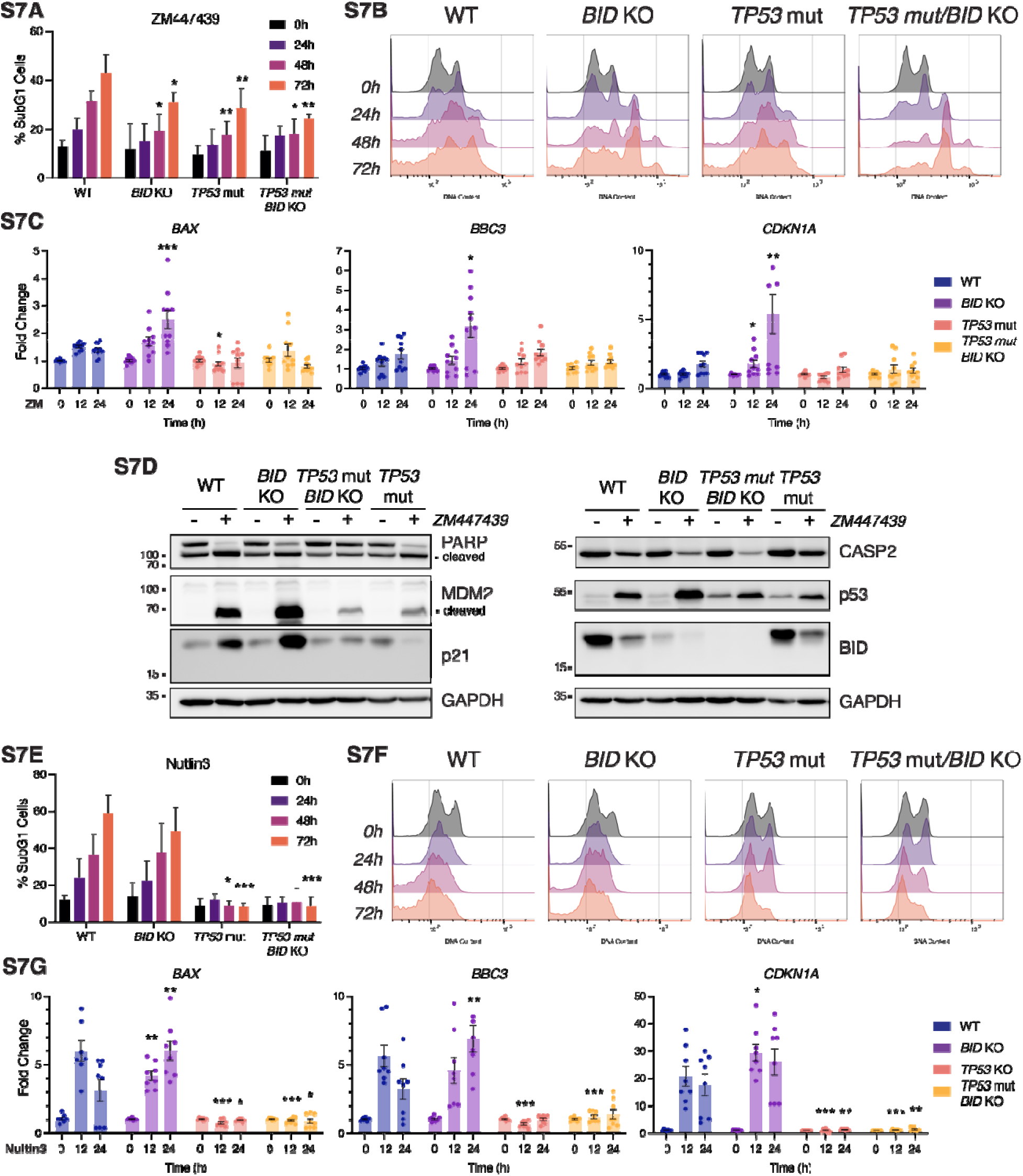
BID and p53 cooperate to induce apoptosis in the p53 proficient Burkitt lymphoma cell line BL2 after cytokinesis failure. **(A)** Quantification of the percentage of subG1 BL2 cells WT and derivative CRISPR/Cas9 edited pools harboring sgRNAs targeting BID, p53 or both at different time points after treatment with 2μM ZM447439. Data is presented as means ± SD of N ≥ 4 independent replicates. Statistics was calculated by one-way ANOVA with Dunnett’s multiple comparison test, comparing each time point of the KO pool to the corresponding time point in the WT sample. * = p value < 0.05; ** = p value < 0.01; ** = p value < 0.001. **(B)** Example of the DNA content profiles of BL2 WT and derivative pools transduced with lentiCRISPR constructs targeting BID, p53 or both in combination at different time points after treatment with 2μM ZM447439. SubG1 events are quantified in Fig. S6C. **(C)** RT-qPCR analysis of the p53 targets BAX, BBC3/PUMA and CDKN1A/p21 on BL2 cells at different time points after 2μM ZM447439 treatment. Results are normalized over the house-keeping gene GAPDH and presented as fold-change over the time point 0h for each polyclonal cell line pool. Data is presented as means ± SEM and individual points represent the value of the independent biological replicate (N = 4). Statistical significance was calculated by one-way ANOVA with Dunnett’s multiple comparison test, comparing each time point of a KO clone to the corresponding time point of the WT sample. * = p value < 0.05; ** = p value < 0.01; *** = p value < 0.001. **(D)** Western blot analysis of BL2 cells and derivative polyclonal pools transduced with lentiCRISPR constructs targeting BID, p53 or both in combination after 48h of treatment with 2μM ZM447439. **(E)** Same as in A but after 10μM Nutlin3 treatment. N ≥ 2 independent biological replicates. **(F)** Same as in B but after 10μM Nutlin3 treatment. **(G)** Same as in C but after 10μM Nutlin3 treatment. N = 3 independent replicates

**Fig. S8:**
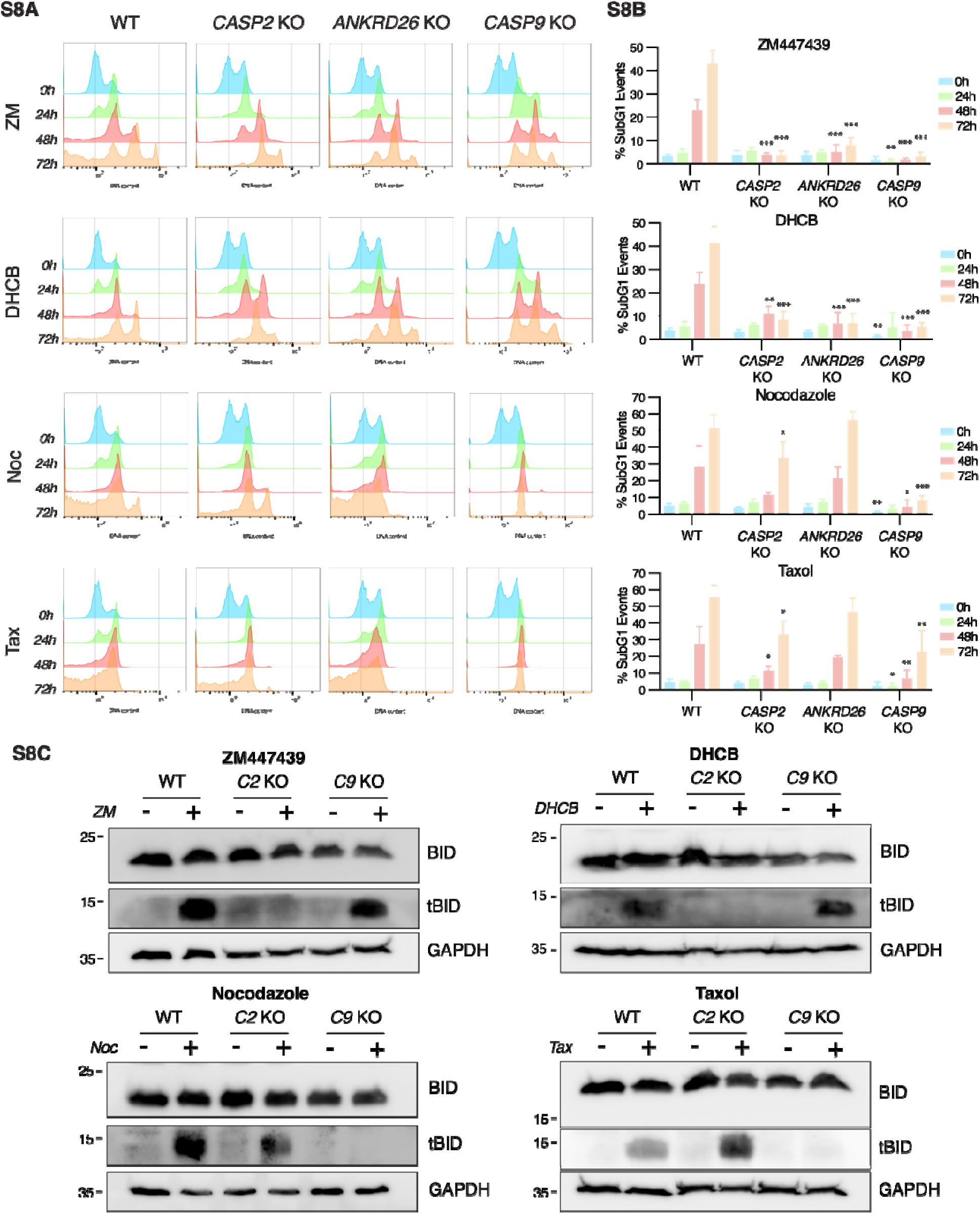
Only cytokinesis failure elicits the centrosome-PIDDosome-tBID axis. **(A)** Representative DNA content profiles of Nalm6 cells WT and derivative clones lacking caspase-2, caspase-9 or ANKRD26 after different time points of exposure to 2μM ZM447439 (ZM), 4μM DHCB, 100nM Nocodazole (Noc) or 50nM Taxol (Tax). **(B)** Quantification of the percentage of subG1 events from B. Data is presented as means ± SD of N ≥ 3 independent biological replicates. Statistical significance was calculated by one-way ANOVA with Dunnett’s multiple comparison test, comparing each time point of the KO clone to the corresponding time point of the WT sample. * = p value < 0.05; ** = p value < 0.01; *** = p value < 0.001. **(C)** Representative western blots showing the processing of BID into tBID in Nalm6 cells WT or clones lacking caspase-2 or caspase-9 after 24h of treatment with the drugs described in A.

**Fig. S9:**
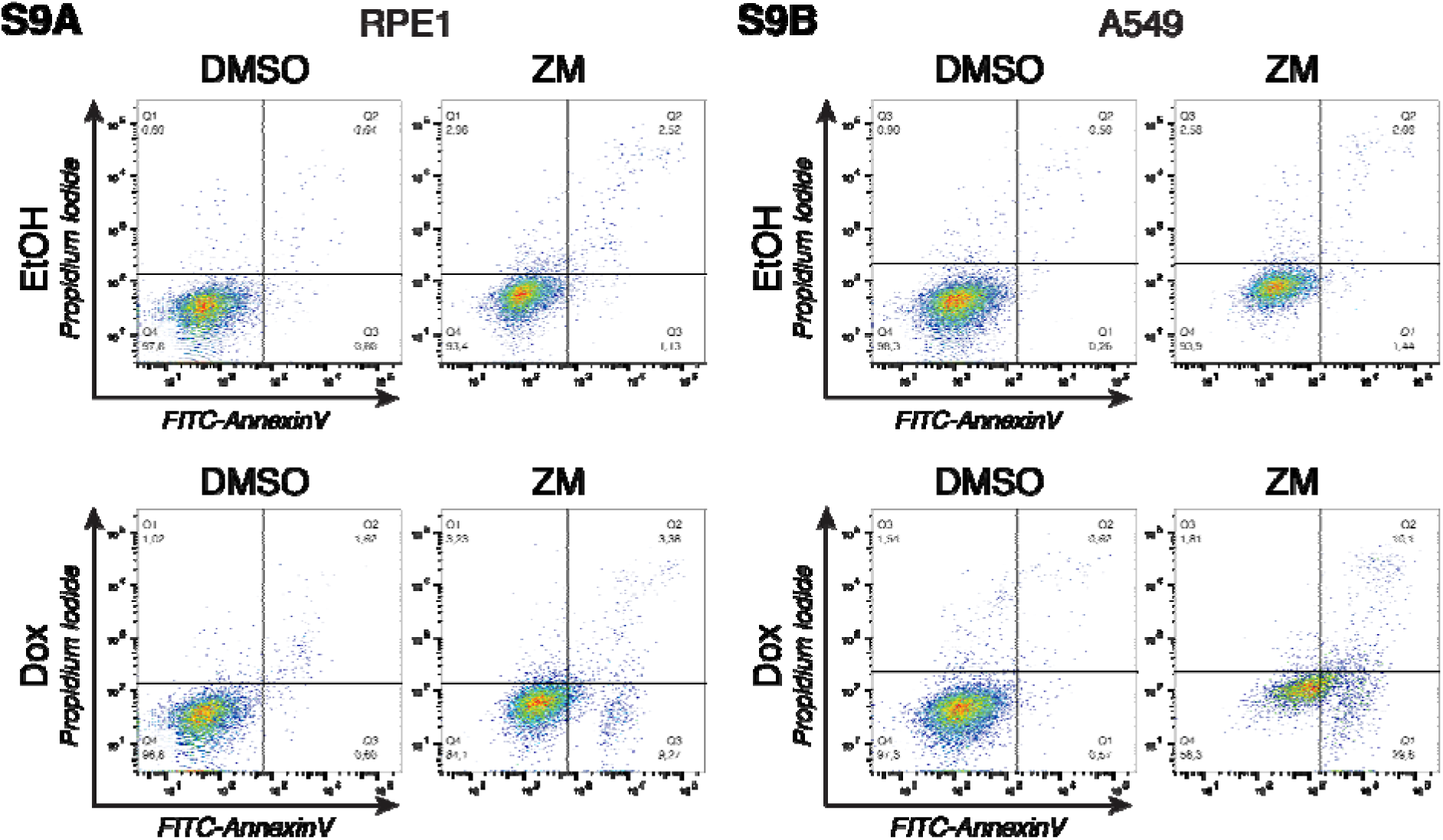
BID overexpression sensitizes epithelial cells to cell death after cytokinesis failure. **(A)** Representativ dot-plot examples of AnnexinV/PI staining followed by flow cytometric analysis of RPE1 cells overexpressing BID after treatment with 2μM ZM447439. **(B)** A549 cells treated the same as in A.

